# Probing neuronal functions with precise and targeted laser ablation in the living cortex

**DOI:** 10.1101/2020.09.09.289041

**Authors:** Zongyue Cheng, Yiyong Han, Bowen Wei, Baoming Li, Meng Cui, Wenbiao Gan

## Abstract

Targeted cell ablation is an important strategy for dissecting the function of individual cells within biological tissues. Here we developed an amplified femtosecond laser-coupled two-photon microscopy (AFL-TPM) system that allows instantaneous and targeted ablation of individual cells and real-time monitoring of neuronal network changes in the living mouse cortex. Through precise and iterative control of the laser power and position, individual cells could be ablated by a single femtosecond light pulse with minimum collateral damage. We further show that ablation of individual somatostatin-expressing interneuron increases the activity of nearby neurons in the primary motor cortex during motor learning. Through precise dendrotomy, we reveal that different dendritic branches of layer 5 pyramidal neurons are structurally and functionally independent. By ablating individual cells and their processes in a spatiotemporally specific manner, the AFL-TPM system could serve as an important means for understanding the functions of cells within the complicated neuronal network.

## Introduction

Cell ablation is an important strategy for understanding the functions of cells within biological tissues(Grégoire and Kmita, 2014; Tertilt et al., 2005). Various chemical-, genetic-, optogenetic- and laser-based methods have been employed for cell ablation(Makhijani et al., 2017). The chemical method uses small molecules to initiate apoptosis and cause a wide range of cell death(Ben-David and Benvenisty, 2014). The genetic-based ablation is typically achieved by activating apoptotic signals in specific cell types by expressing toxins or toxin receptors under the control of cell-type specific promoters(Hu et al., 2008; Saito et al., 2001). This method has been used successfully for dissecting the functions of cells in the immune system(Ito et al., 2017) and the nervous system(Cheng et al., 2016; Ralvenius et al., 2015). Furthermore, the optogenetic method has been applied to induce apoptotic responses of specific cell types by generating excessive reactive oxygen species after light irradiation of cells expressing phototoxic fluorescent proteins(Kobayashi et al., 2013; Makhijani et al., 2017). More recently, a combination of light-irradiation and chemical methods have been shown to damage the DNA of individual cells by bleaching a nucleic acid-binding dye *in vivo*(Damisah et al., 2017). In addition to the methods of inducing cell apoptosis, laser-based ablation can cause necrotic cell death through rapid accumulation of photothermal energy in targeted individual cells(Makhijani et al., 2017; Tsai et al., 2009; Vogel et al., 2005) in *Caenorhabditis elegans*(Churgin et al., 2013; Fouad et al., 2018), zebrafish(Naylor et al., 2018, 2016), and hair follicles(Beiko et al., 2011; Rompolas et al., 2012).

Specificity, accuracy, and effectiveness are critical factors to consider in practical applications of targeted cell ablation(Wang et al., 2013). In this regard, each of the current cell-ablation methods has its pros and cons. For example, the chemical-based method may lack ablation specificity, while genetic or optogenetic ablation requires the expression of toxins in specific cell types, which is often challenging and difficult to achieve. In addition, all the chemical-, genetic-, optogenetic-based methods involve cell apoptosis, which takes place over a relatively long period of time (hours to days) and may lead to compensatory responses of biological tissues(Dunn, 2015a). Although the typical pulse laser-based method can cause rapid cell ablation(Allegra Mascaro et al., 2013; Eyo et al., 2014), the limited energy generated by the Ti:sapphire oscillator pulses (∼ 1-10 nJ per single femtosecond pulse) requires irradiating cells for a period of seconds to ensure a thorough deactivation of targeted cells(Hayes et al., 2014; Orger et al., 2008; Pozner et al., 2015). As such, this process would often cause collateral damages to the adjacent tissues because of the cumulative thermal effects(Bauer et al., 2015; Di Niso et al., 2014; Jasiński, 2018). Moreover, although Ti:sapphire lasers have been reported to ablate cells on the cortical surface(Buffelli et al., 2007; Park et al., 2019), the limited energy of each laser pulse makes it difficult to ablate cells located deep in the brain.

The amplified femtosecond laser generated by chirped pulse amplification (CPA) technology can achieve a high pulse power (∼0.3 mJ) with a low repetition rate (up to tens of kHz)(Krüger and Kautek, 2012; Strickland and Mourou, 1985) and ablate targeted cells rapidly without linear heat accumulation(Tsai et al., 2009).With those advantages, this amplified femtosecond laser has been performed to cut the axons of worms(Bourgeois and Ben-Yakar, 2008; Chung and Mazur, 2009; Gabel et al., 2008; Guo et al., 2008; Morris et al., 2013; Wu et al., 2007; Yanik et al., 2004), model stroke by insulting the subsurface cortical vasculatures(Blinder et al., 2012; Nishimura et al., 2007, 2006; Tsai et al., 2003), and ablate cells in cultures(Hamad, 2016) or brain slices(Tsai et al., 2009). However, due to the lacking of precise power and position control, the amplified femtosecond laser has not been demonstrated for ablating individual neurons and their processes in the living brain.

In this study, we developed an amplified femtosecond laser-coupled two-photon microscope (AFL-TPM) system, which can target any individual cells within a three-dimensional volume with a single laser pulse by precise and iterative control of the laser power and position. The AFL-TPM system enables us to instantaneously and precisely ablate mouse cortical cells *in vivo* and to monitor subsequent changes in the neural network in a real-time fashion. In combination with two-photon imaging of neuronal structure and functions, the AFL-TPM system described here can serve as a promising platform for understanding the roles of different cell populations in the complex neural network.

## Results

### Design of the amplified femtosecond laser ablation system coupled with two-photon laser scanning microscope

To achieve efficient and targeted ablation of brain cells, we configured a regenerative amplifier to generate 35 fs 800 nm laser pulses with high peak powers (up to 0.3 mJ per pulse). The 10 kHz clock signal from the amplifier was used to trigger a Pockels cell, which selectively deliver a single pulse for targeted ablation. The amplifier beam was combined with the imaging beam (140 fs, 930 nm, 80 MHz) before entering the laser scanning microscope (**Fig. 1a,b, Supplementary Fig. 1a**). By recording the two-photon excited signal from 0.5 μm diameter fluorescence beads, we quantified the focus profile of the 800 nm ablation beam, which shows 0.55, 0.64, and 3.2 μm in *x, y*, and *z* direction (**Fig. 1c**). Moreover, we utilized a series of single pulses (∼0.05 μJ) of 800 nm ablation beam to bleach a fluorescence reference slide, which showed darkened regions of submicron in size (**Fig.1d,e**).

**Figure 1.**
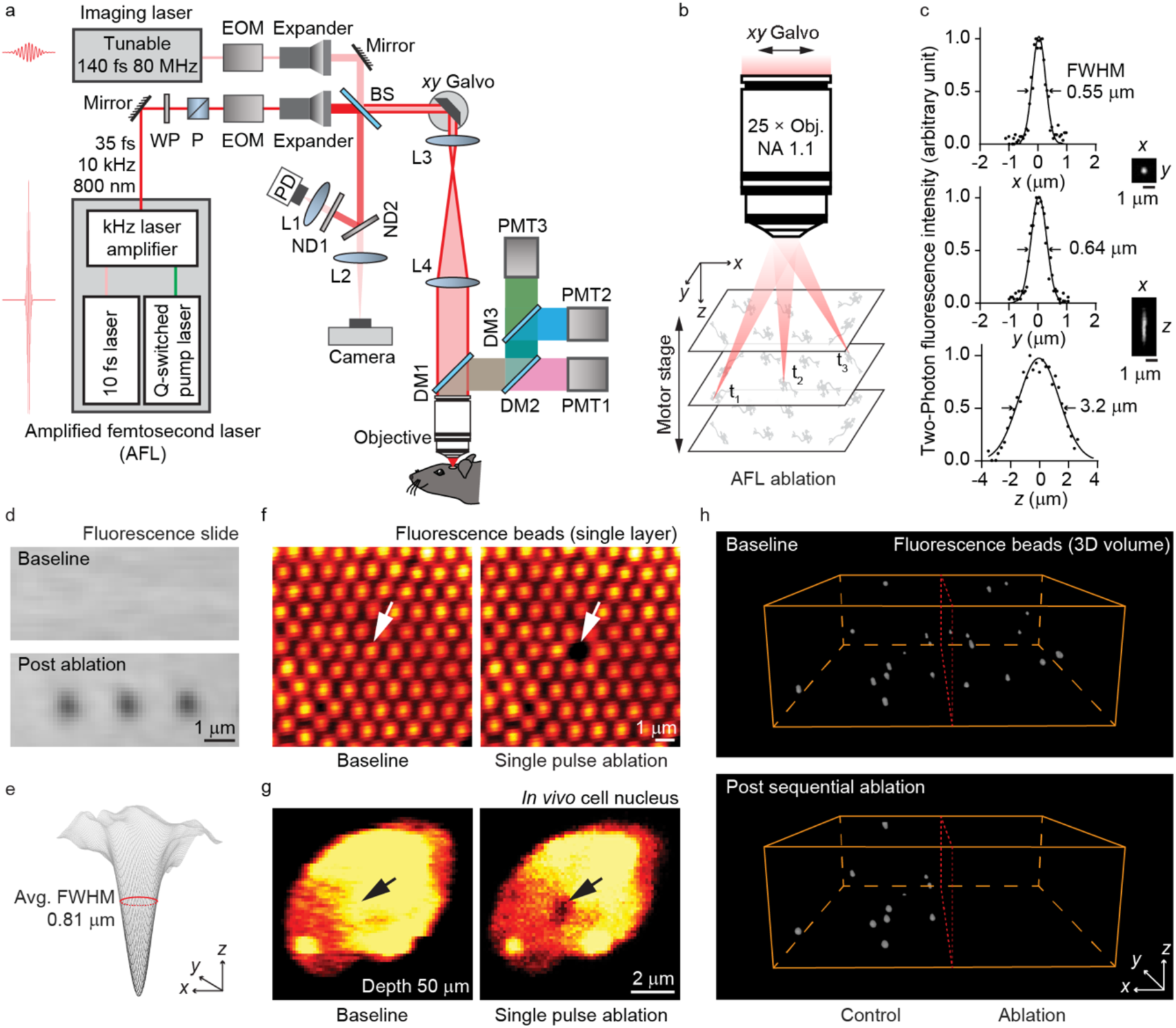
Target ablation at submicron resolution with AFL-TPM. **a**, Schematic of modified two-photon microscope coupled with amplified femtosecond laser and tunable imaging scanning laser. WP, waveplate; P, polarizer; EOM, electro-optic modulators; BS, beam splitter; PD, photodiode; ND, neutral density filter; DM, dichroic mirror; PMT, photomultiplier tube; L, lens. **b**. Schematic of AFL-TPM volume ablation by *x*y Galvo and motor stage. t_1_ to t_3_ indicates ablation at different instants of time. **c**, Measured point spread function (PSF) of 800 nm amplified femtosecond laser with ∼ 500 nm fluorescence bead. Maximum intensity projection (MIP) of the bead images in *xy and xz* directions are shown on the right. With Gaussian fitting, the full width at half maximum (FWHM) is 0.55, 0.64, and 3.2 μm in *x, y*, and *z* direction. **d**, The AFL-TPM-induced ablation (0.05 μJ in a single pulse) on a fluorescence reference slide at the submicron level. Black holes indicate the targeted points, scale bar: 1 μm. **e**, Averaged FWHM of irradiated points in **d** is 0.81 ± 0.03 μm. **f**, Representative images showing AFL-TPM system-mediated (0.71 μJ in a single pulse) precise ablation of single layer fluorescence beads. White arrows point to the targeted bead. Scale bar: 1 μm. **g**, Representative images showing AFL-TPM system-mediated precise ablation of the cell nucleus *in vivo*. Hoechst 33342 was used to label the nuclei of live cells *in vivo*. A black spot was observed right after a single pulse (0.21 μJ) irradiated on the targeted nucleus. Images depth: 50 μm, scale bar: 2 μm. **h**, 3D volume (*x, y*: 46 μm, Z: 50 μm) ablation of fluorescence beads. The targeted fluorescence beads (∼1 μm) embedded in 2.2% agarose were sequentially irradiated with 0.09 μJ pulses.

A key requirement for subcellular ablation in the living brain is the focus positioning accuracy. To accomplish this, we developed an iterative convergence method based on the slow-axis Galvo position sensor feedback to accurately control the ablation position (see Methods). For the accuracy test, we prepared a single layer of 1 μm fluorescence beads on a cover glass (**Fig.1f**, left) and selectively targeted an individual bead. After a 0.71 μJ single pulse ablation, only the targeted bead was bleached and none of the surroundings beads was affected (**Fig. 1f**, right). In addition, we also targeted the nucleus of fluorescently-labeled neuronal cells with a 0.21 μJ single pulse in the living mouse cortex. Similar to the results of targeted beads, the targeted nucleus also showed a darkened region in the living mouse cortex (**Fig.1g**). For ablation within 3D volume, we combined the transverse positioning of Galvo and the axial positioning of a motorized high precision stage. After the user selection of the beads location within the 3D image stack, the control program could automatically apply individual laser pulses to target beads at each location (**Fig. 1h, Supplementary Movie 1**).

Together, these results demonstrate that our amplified femtosecond laser-coupled two-photon microscope (AFL-TPM) system can rapidly carry out high-precision targeted ablation within a 3D volume.

### Targeted ablation of cells deep in the cortex

To evaluate the accuracy and efficiency of AFL-TPM system-mediated cell ablation, we applied two commonly used nucleus staining dyes, Propidium Iodide (PI) and Hoechst 33342, to the mouse cortex in order to label dead and live cells, respectively (**Fig. 2a**). Before the AFL-TPM irradiation, the vast majority of the cells were positive for Hoechst (shown as cyan), with very few being labeled by PI (red) (**Fig. 2b**). However, after the targeted cell in the depth of 20-40 μm was irradiated by a single pulse that ranged from 0.13 to 0.16 μJ, the Hoechst signal was reduced to ∼63% of its original value within 1 min **(Supplementary Movie 2)** and further dropped to 50% after 30 mins (**Fig. 2b,d**). On the other hand, the intensity of PI in the targeted cells increased immediately after laser irradiation (**Fig. 2b,d, Supplementary Movie 2**). No significant changes in the Hoechst and PI intensity were observed in the surrounding untargeted cells (**Fig. 2b,c**). These results show rapid damage of cell nuclei after a single laser pulse with the AFL-TPM system.

**Figure 2.**
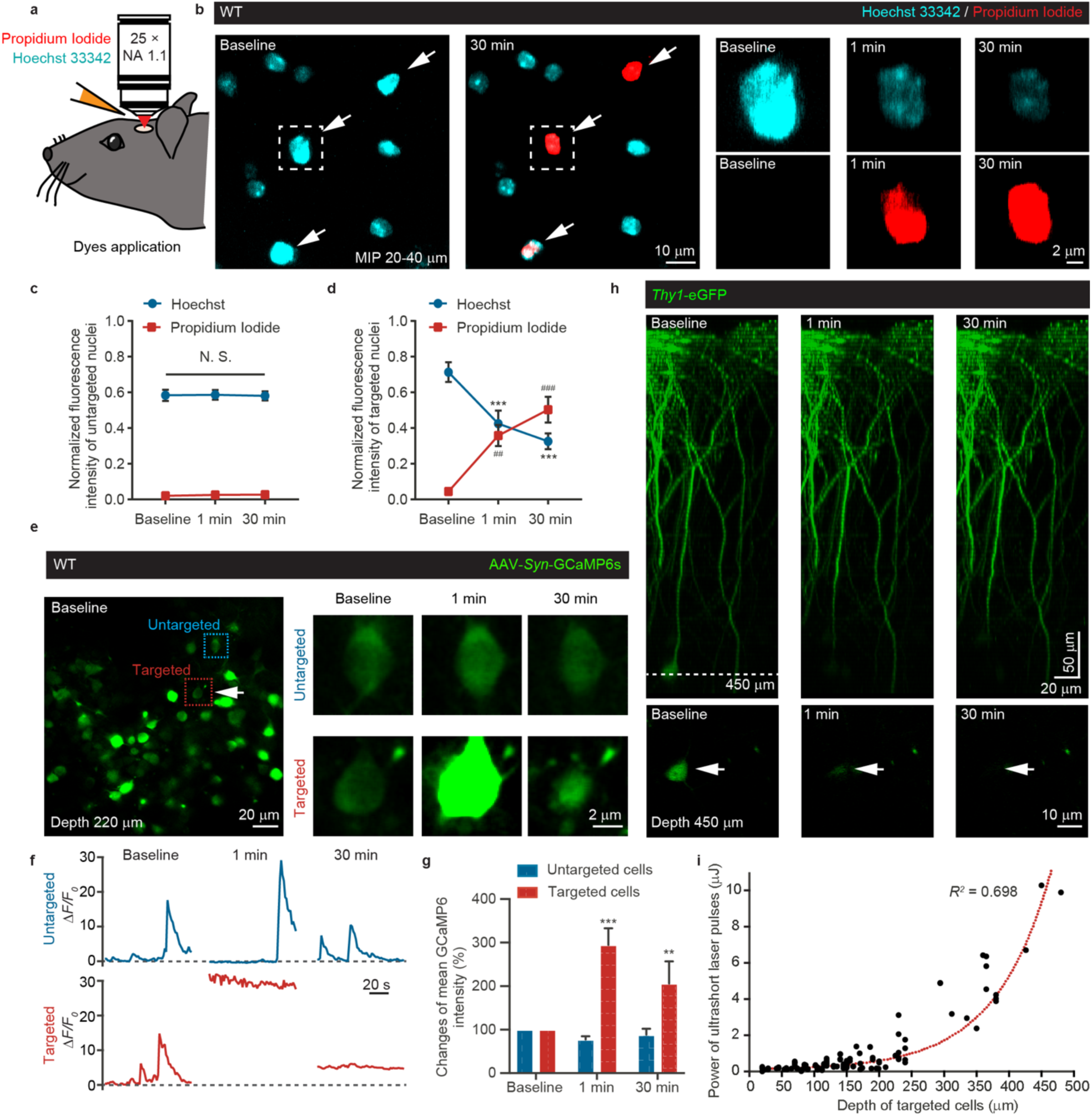
AFL-TPM system-mediated ablation of neuronal cells in the living mouse brain. **a**, Schematic of *in vivo* labeling of live and dead cell fluorescent dyes before two-photon imaging. Hoechst 33342 (cyan, 0.04 mg/ml) and Propidium Iodide (red, 0.5 mg/ml) were applied to the mouse cortex for 15 min before imaging. **b**, Two-photon images showing the nuclei of live cells (cyan) and dead cells (red) before (baseline) and after (1 min, 30 min) AFL-TPM system-mediated ablation (0.13-0.16 μJ). Arrows point to targeted cells. The targeted cell in the white dotted boxes showed time-dependent changes of fluorescent labeling at a higher magnification (right panels). Images depth: 20-40 μm. Scale bars: 10 μm (left) and 2 μm (right). **c**, Fluorescence intensities of Hoechst and PI showing no changes in the nearby untargeted cells after the targeted cell was ablated by the AFL-TPM. *n =* 64 cells from 3 mice. Dunnett’s multiple comparisons test. N.S., not significant. **d**, A time-dependent increase in PI fluorescence (red, baseline vs. 1 min, *P* = 0.0155; baseline vs. 30 min, *P* < 0.0001) and decrease in Hoechst fluorescence (cyan, baseline vs. 1 min, *P* < 0.0001; baseline vs. 30 min, *P* < 0.0001) in AFL-TPM irradiated cells. *n =* 14 cells from 3 mice. Dunnett’s multiple comparisons test. **e**, Representative *in vivo* time-lapse images showing calcium signals (AAV-*Syn*-GCaMP6s, green) in the targeted neuron (arrow) and its nearby neurons at baseline and after AFL-TPM irradiation (1.25 μJ). The calcium signal decreased in the targeted cell over time but remained significantly higher than that of their nearby cells 30 min post laser irradiation. Images depth: 220 μm, scale bars: 20 μm (left) and 2 μm (right). **f**, The targeted cell (red dotted box in **e**) but not the nearby cell (blue dotted box in **e**) showing blunted spontaneous calcium activity post AFL-TPM irradiation. *F*_*0*_ of baseline was used during the Δ*F/F*_*0*_ quantification of 1 min and 30 min. **g**, Relative GCaMP6 intensity changes in the targeted cells and their nearby cells post AFL-TPM irradiation. A significant increase in calcium concentration as measured with GCaMP6 was observed in targeted cells (baseline vs. 1 min, *P* < 0.0001; baseline vs. 30 min, *P* = 0.0017) but not in the nearby cells. *n =* 15 targeted cells and 17 nearby (untargeted) cells from 5 mice. Dunnett’s multiple comparisons test. **h**, The AFL-TPM efficiently ablated mouse neuronal cells in deep brain tissue (white dotted line) without compromising the morphological structures of the surface region. The fluorescence intensity of *Thy1*-eGFP labeled layer 5 pyramidal soma (arrows, bottom) reduced rapidly after AFL-TPM irradiation (10.28 μJ). Top: 3D reconstruction from −10 to 470 μm of the brain tissues, scale bars: 20 μm (*xy*) and 50 μm (*z*); bottom: single-slice images at 450 μm, scale bar: 10 μm. **i**, The pulse energy required for effective AFL-TPM ablation was positively correlated with tissue depth. *n =* 123 cells from 55 mice, *R*^*2*^ = 0.698. Data are mean ± s.e.m. **d, g**, **or^##^ *P* < 0.01, ***or ^###^ *P* < 0.001.

To further investigate changes of targeted neurons induced by laser irradiation, we performed calcium imaging of layer 2/3 neurons expressing the genetically encoded calcium indicator GCaMP6 in the motor cortex before and after the ablation. We found that GCaMP6 signals in the targeted neurons were 3 times higher than the baseline level 1 min after laser irradiation and remained 2 times higher at 30 mins (**Fig. 2e,g, Supplementary Movie 3**). Notably, spontaneous calcium transients were not observed in the targeted cell after ablation, but were frequent in the surrounding untargeted cells (**Fig. 2f, Supplementary Movie 3**). Furthermore, the targeted neurons showed no changes of somatic calcium in response to treadmill running (**Supplementary Fig. 2 a-d**). In addition to ablating layer 2/3 neurons, the near infrared wavelength, high-order nonlinearity and the high pulse energy generated by the amplified femtosecond laser allowed us to ablate cells embedded deep in the brain tissue. The laser power required for ablating targeted cells was positively correlated with the depth of the cells within the tissue (**Fig. 2i**). At 10.28 μJ power of single pulse, we successfully ablated targeted pyramidal neurons in cortical layer 5 without damaging the superficial layer of the cortex in *Thy1*-eGFP M line transgenic mice (**Fig. 2h**).

Collectively, the above observations demonstrate that a single pulse generated by the amplified femtosecond laser is sufficient to ablate targeted cells located in different cortical layers.

### The AFL-TPM system allows the ablation of targeted cells at the subcellular resolution

To determine the potential collateral damages associated with the targeted cell ablation by AFL-TPM, we examined the morphological changes of both AFL-TPM ablated neurons and its surrounding tissues in *Thy1*-eGFP mice. Within 1 min after laser irradiation, we found that neuronal somata swelled while the fluorescence of dendrites and axons of the targeted neuron diminished, which may owe to the broken of cytomembrane. The gross structure of the targeted neurons was invisible 30 min after laser irradiation. In contrast, there was no detectable structural change in neighboring untargeted neurons few microns away (**Fig. 3a-c, Supplementary Movie 4**).

**Figure 3.**
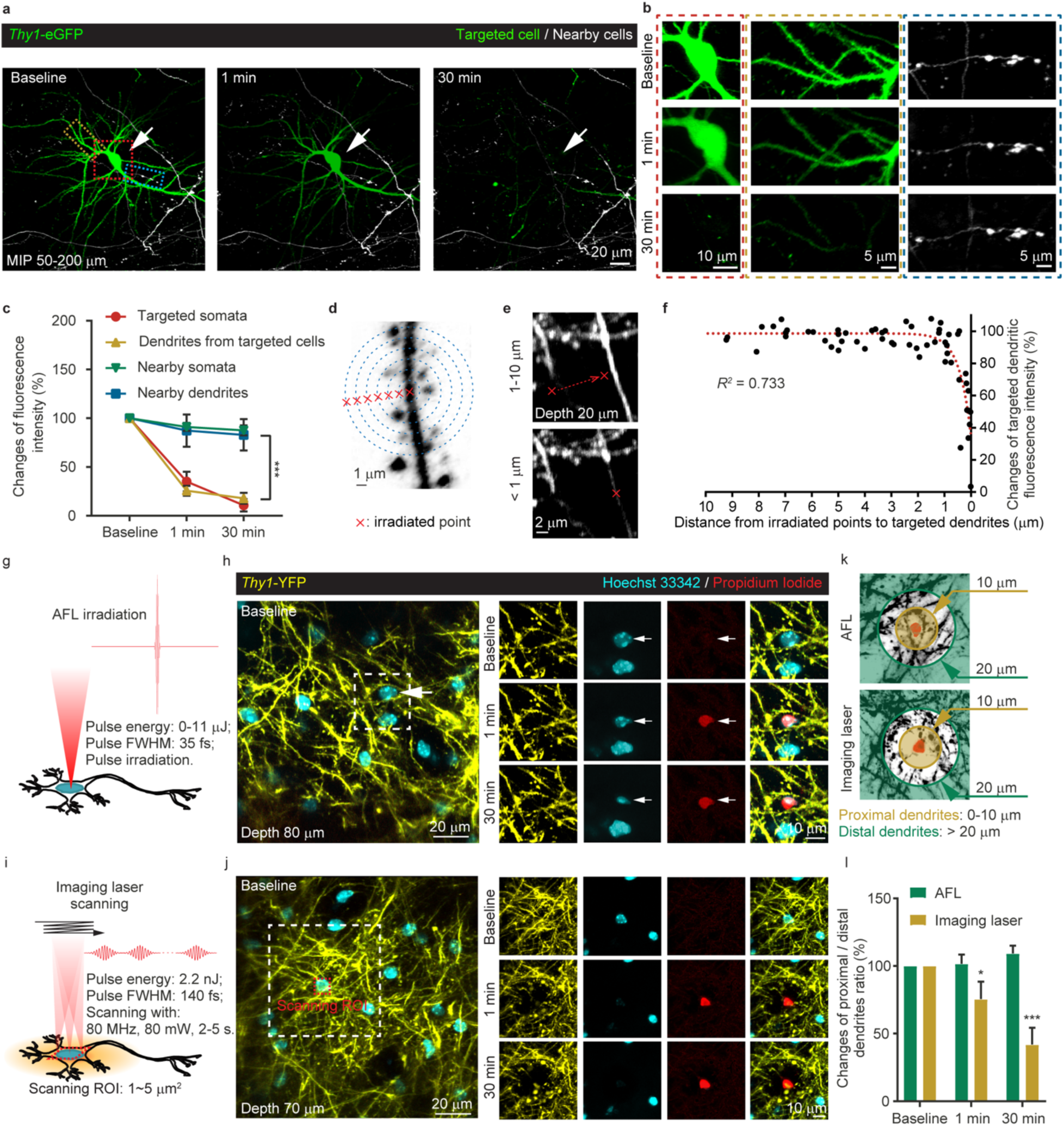
Precise and targeted ablation of cells at the subcellular level with AFL-TPM. **a**, The fluorescence signal of the targeted neuron (arrow, red dotted box) was reduced 1 min after AFL-TPM irradiation (0.71 μJ) and completely diminished 30 min later. The structural integrity of the nearby cells (white) was unaffected. Images depth: 50-200 μm. Scale bar: 20 μm. **b**, High magnification images of the three color-coded dotted boxes in **a**. The red dotted box shows the targeted soma, yellow shows the dendrites from the targeted soma, and blue shows axons of a nearby cell. Scale bars: 10 and 5 μm. **c**, Changes of fluorescence intensity of the AFL-TPM targeted cells and their nearby cells. *n =* 10, 4 somata and 16, 15 dendrites from 8 mice. Dunnett’s multiple comparisons test. **d**, Schematic diagram of the relationship between AFL-TPM irradiation distance from the targets and effectiveness of ablation of the targeted dendrites. Red cross symbols denote different irradiated points with the same pulse energy. The blue circles are used as references for the distances between the irradiation points and targeted dendrites. Scale bar: 1 μm. **e**, Images showing the irradiation distance needs to be within 1 μm to achieve effective ablation of the targeted dendrites. Red cross symbols denote the irradiation spots (0.14 μJ). Scale bar: 1 μm. **f**, A nonlinear logarithmic curve showing the relationship between the AFL-TPM irradiation distance and the effectiveness of ablation of the targeted dendrites. The red dotted line is the fitting curve, *R*^*2*^ = 0.733. Each black dot represents a distance of the irradiation point and the targeted dendrite against the relative fluorescence intensity changes in the targeted dendrite after each AFL-TPM ablation. *n =* 63 ablations from 3 mice. **g**, Schematic of AFL-TPM system-mediated cell ablation with a single pulse. The pulse energy was confined between 0-11 μJ with a fixed duration of 35 fs. The waveform of the laser pulse is displayed at the upper-right corner of the panel. **h**, The AFL-TPM ablated (0.14 μJ) the targeted cell (arrows) without affecting the viability of nearby neurons (cyan) and structural integrity of dendrites (yellow). Time-lapse fluorescence intensity of the AFL-TPM targeted cell and its nearby tissue in the white dotted box are displayed on the right panels. A successful cell ablation was recognized by a significant increase in PI fluorescence and a decrease in Hoechst fluorescence. Images depth: 80 μm. Scale bars: 20 μm (left) and 10 μm (right). **i**, Schematic of two-photon laser scanning-mediated cell ablation with multiple pulses. The two-photon laser scanned a small nucleus region (1∼5 μm^2^) of interest for 2-5 s with the average power of 80 mW. The energy for each pulse was 2.2 nJ. The orange area denotes the thermal effect generated by line scanning accumulation. **j**, The low pulse power imaging laser scanning damaged neuronal structures surrounding the targeted cell due to heat accumulation (80 mW, 2 s). Images were taken from the same mouse in **h** but in a different area. Images depth: 70 μm, scale bars: 20 μm (left) and 10 μm (right). **k**, Diagrams showing dendrites at the border and remote regions of targeted cells. The light-yellow area denotes the proximal dendrites within 10 μm diameter of the targeted cell and the light-green area denotes the distal dendrites that are more than 20 μm away from the targeted cell. **l**, Relative ratio changes of the fluorescence intensity of proximal to distal dendrites at different time points (baseline vs. 1 min, *P* = 0.0432; baseline vs. 30 min, *P* < 0.0001). *n =* 18 cells for the amplified femtosecond laser group and 9 for the imaging laser group from 7 mice. Dunnett’s multiple comparisons test. Data are mean ± s.e.m. **c, l**, * *P* < 0.05, *** *P* < 0.001.

To further determine the accuracy and confinement of the AFL-TPM system-mediated cell ablation *in vivo*, we focused the laser at varying distances to layer 1 dendrite of *Thy1*-eGFP mice (**Fig. 3d**). We found that the effective distance for the AFL-TPM to ablate the targeted dendrites was within 1 μm (**Fig. 3e,f**). When the same measurement was performed on blood vessels, we observed that vascular damage was detectable only when the irradiation point was set on the capillary (**Supplementary Fig. 3c**).

The above observations suggest that unlike the Ti:sapphire laser-based ablation(Hayes et al., 2014; Orger et al., 2008; Pozner et al., 2015), an individual pulse of amplified femtosecond laser does not cause a localized heat accumulation effect during cell ablation and therefore causes confined damage. To investigate this issue further, we ablated individual neurons with either the AFL-TPM or the Ti:sapphire laser and subsequently compared potential damage to nearby dendrites in *Thy1*-YFP mice (**Fig. 3h,j**, shown in yellow). In this experiment, the mouse cortex was stained with PI and Hoechst prior to laser irradiation. We observed that the AFL-TPM method damaged the nucleus of an individual neuron while leaving the surrounding dendrites undamaged (**Fig.3g,h**). We observed that the AFL-TPM method damaged the nucleus of an individual neuron while leaving the surrounding dendrites undamaged (**Fig.3g,h**). Moreover, the blood vessels around the targeted structure also remained intact (within 1 mm of the target, **Supplementary Fig. 3a,b**). For comparison, we utilized the 80 MHz repetition rate imaging beam (two-photon Ti:sapphire oscillator) to ablate individual neurons by scanning a small area that is commonly used in the brain damage model(Buffelli et al., 2007; Fu et al., 2014; Hayes et al., 2012; Li et al., 2018; Pozner et al., 2015; Rompolas et al., 2012) and found significantly decreased YFP intensity and compromised structural integrity in the area surrounding the ablation site. We also parked the laser beam in single point with different time and found the similar result (**Fig. 3i,j; Supplementary Fig. 4**). When the ratio of fluorescence intensity of proximal dendrites (border region, at a radial distance < 10 mm from the targeted cell) to distal dendrites (remote region, radial distance > 20 mm) was compared between both groups (**Fig. 3k**), we found that the ratio did not change in the amplified femtosecond laser ablation group over the 30-min observation period, but reduced by 25% and 40% at 1 min and 30 min respectively, after irradiation with the Ti:sapphire imaging laser (**Fig. 3l**).

Together, these results highlight that the AFL-TPM is superior to the commonly used two-photon scanning femtosecond pulsed laser in achieving precise cell ablation without incurring collateral damages to the surrounding tissues at the subcellular level.

### The AFL-TPM system-mediated ablation of somatostatin-expressing interneurons enhances the excitability of nearby neurons during motor training

The precisely targeted cell ablation with the AFL-TPM system allows us to manipulate cells of interest at a confined spatial scale to determine their functional roles *in vivo*. Previous studies have demonstrated that somatostatin (SST)-expressing interneurons extend their axonal projections onto layer 2/3 pyramidal neurons(Adesnik et al., 2012; Fino and Yuste, 2011; Pfeffer et al., 2013; Urban-Ciecko and Barth, 2016; Xu et al., 2013). Inhibition of SST interneuron population directly with an optogenetic method(Gentet et al., 2012) or indirectly through the activation of VIP interneurons(Fu et al., 2014) enhanced the activity of pyramidal neurons. Recent studies have also revealed that SST interneurons were critical for inducing and maintaining the sequential activation of layer 2/3 pyramidal neurons in mice during running by CNO-induced GPCR regulation(Adler et al., 2019). To determine the impact of individual layer 2/3 SST interneurons on their surrounding neurons *in vivo*, we expressed cre-dependent tdTomato in SST interneurons and GCaMP6 in both pyramidal neurons and interneurons in layer 2/3 of the primary motor cortex of SST-Cre mice (**Fig. 4a,b**). We used calcium imaging to monitor treadmill running-induced activity changes in surrounding neurons in the primary motor cortex with two-photon microscope before and after targeted SST interneurons were irradiated by the amplified femtosecond laser (**Fig. 4a**).

**Figure 4.**
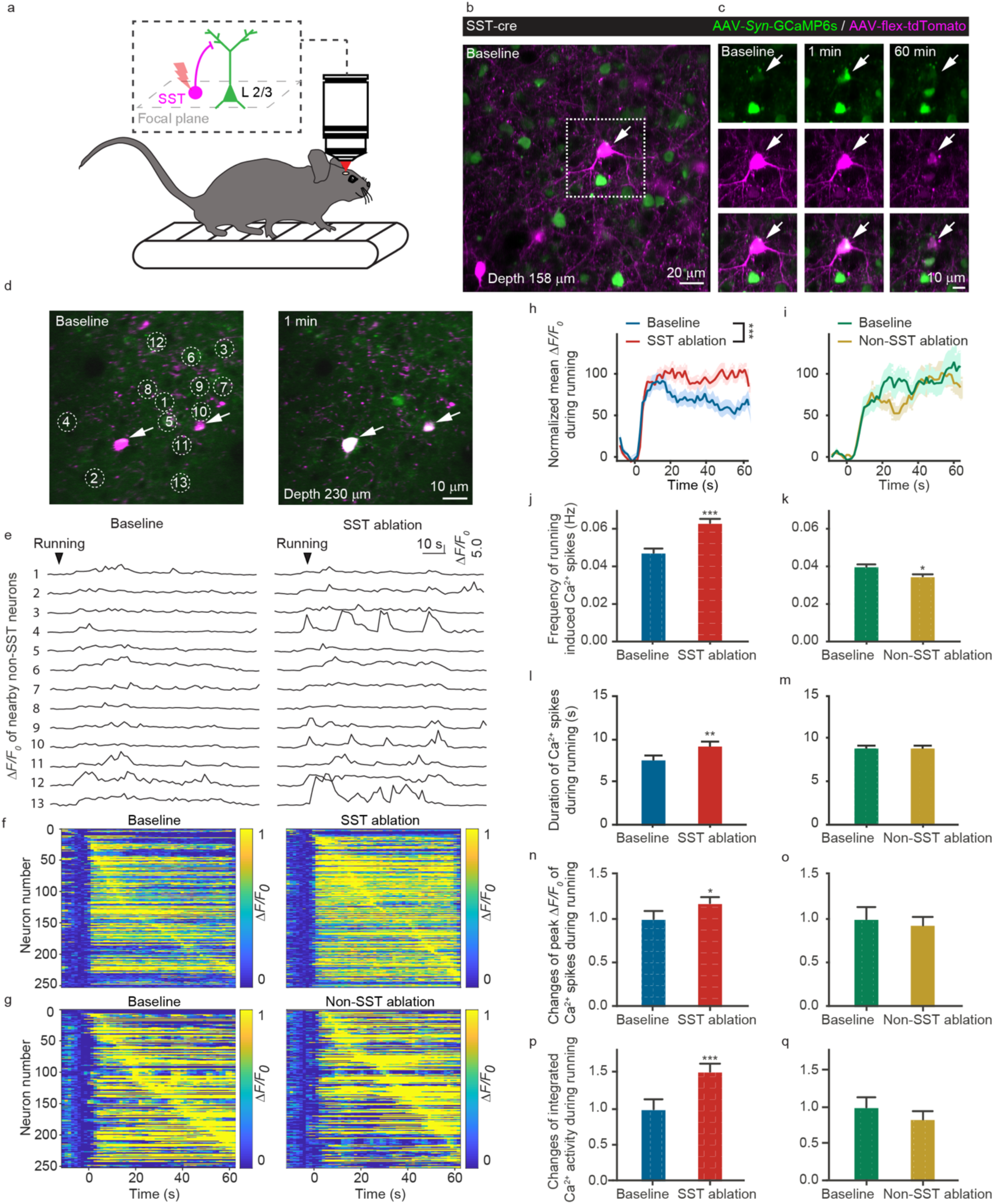
AFL-TPM-mediated ablation of somatostatin (SST)-expressing interneurons increases activity of nearby neurons. **a**, Schematic of *in vivo* two-photon calcium imaging of layer 2/3 SST interneurons and non-SST neurons when the head-restrained mouse was running forward on a treadmill. **b,c**. Representative images showing that AFL-TPM can precisely ablate the targeted SST interneurons (arrows). Two-photon imaging showing the changes in fluorescence intensity of flex-tdTomato (magenta) labeled SST interneuron and *Syn*-GCaMP6s labeled calcium signals (green) before and after laser irradiation (0.48 μJ) (**b, c**). Both fluorescence signals were diminished to the negligible levels 60 min post ablation. Images depth: 158 μm. scale bars: 20 μm (left) and 10 μm (right). **d**, *In vivo* imaging showing effective ablation of 2 SST interneurons (arrows) in layer 2/3 of the primary motor cortex by AFL-TPM (0.63-0.74 μJ). Images depth: 230 μm, scale bar: 20 μm. **e**, Representative running-induced calcium traces of nearby non-SST neurons from awake, head-restrained mice before (left) and after (right) ablation of the targeted SST interneurons in **d**. The black arrow denotes the start point of running. **f,g**, Heatmaps showing the average activity of nearby neurons (rows) in response to forward running before (left) and after (right) AFL-TPM-mediated ablation of SST interneurons (**f**) or non-SST neurons (**g**). Time 0: treadmill on. Neurons were ordered according to the time of max peak Δ*F/F*_*0*_ responses. The value of all calcium signals between 0-1 was arranged from blue to yellow. *n =* 252 cells from 6, and 4 mice. **h,i**, Population average of running-induced calcium responses were significantly increased (*P* < 0.0001) in the SST ablation group (**h**) but not in the non-SST ablation group (**i**). Tint colors: ± s.e.m. *n* = 252 cells from 6, and 4 mice. Kolmogorov-Smirnov test. **j,k**, The frequency of running-induced calcium transients (peak Δ*F/F*_*0*_ > 0.5) was significantly increased in the SST ablation group (**j**, *P* < 0.0001 vs. baseline. *N =* 166, 177 cells from 6 mice. Mann Whitney test) but decreased in the non-SST ablation group (**k**, *P* = 0.0376 vs. baseline. *n =* 193, 201 cells from 4 mice. Mann Whitney test). **l,m**, The duration of running-induced calcium spikes increased in the SST ablation group (**l**, *P* = 0.0062 vs. baseline. *n =* 166, 177 cells from 6 mice. Mann Whitney test) but not in the non-SST ablation group (**m**, *P* = 0. 9266 vs. baseline. *n =* 193, 201 cells from 4 mice. Mann Whitney test). **n,o**, The peak Δ*F/F*_*0*_ of running-induced calcium spikes increased in the SST ablation group (**n**, *P* = 0.0108 vs. baseline. *n =* 166, 177 cells from 6 mice. Mann Whitney test) but not in the non-SST ablation group (**o**, *P* = 0. 4743 vs. baseline. *n =* 193, 201 cells from 4 mice. Mann Whitney test). **p,q**, Running-induced integrated calcium activity increased in the SST ablation group (**l**, *P* < 0.0001 vs. baseline. *n =* 166, 177 cells from 6 mice. Mann Whitney test) but not in the non-SST ablation group (**m**, *P* = 0.4258 vs. baseline. *n =* 193, 201 cells from 4 mice, Mann Whitney test). **h, j, k, l, n, p**, Data are mean ± s.e.m. * *P* < 0.05, ** *P* < 0.01, *** *P* < 0.001.

Immediately after laser irradiation, targeted SST interneurons showed a burst of calcium transients with little variation of tdTomato fluorescence. Both tdTomato and GCaMP6 fluorescence signals decayed to an undetectable level 30 minutes later, indicating permanent damage of SST interneurons (**Fig. 4c**). When several SST interneurons (approximately 1-3 cells of each view) located in the primary motor cortex were ablated (**Fig. 4d**), we found that the activity of surrounding non-SST neurons (mostly pyramidal neurons) adjacent to irradiated SST interneurons was significantly increased as compared to their activity before ablation (**Fig. 4e,f,h**). Notably, such an increase in calcium signal was absent in neurons when nearby non-SST cells were ablated (**Fig. 4g,i**). We quantified treadmill running-induced calcium transient frequency, duration, variation of peak Δ*F/F*_*0*_, and integrated calcium activity in the surrounding neurons before and after laser irradiation of SST or non-SST neurons. All the four parameters of surrounding non-SST neurons were significantly increased after laser irradiation of SST interneurons (**Fig. 4j,l,n,p**). In contrast, the frequency of calcium transients was decreased in non-SST neurons when the surrounding non-SST cells were ablated (**Fig. 4k**). No difference was detected in the duration, peak Δ*F/F*_*0*_, and integrated calcium activity before and after cell ablation in the same group (**Fig. 4m,o,q**). Additionally, none of the above parameters was affected before and after the ablation of neighboring SST or non-SST neurons when mice were stationary (**Supplementary Fig. 5**).

Together, using the AFL-TPM system-mediated targeted cell ablation method, our results show that only a few of SST-expressing interneurons could suppress the activity of their surrounding neurons as well as the excitability of the local neural networks during motor skill learning.

### Dendrotomy with the AFL-TPM reveals structural and functional independence of pyramidal dendritic branches

It has been suggested that individual dendritic branches act as an independent unit for input integration and synaptic plasticity(Cichon and Gan, 2015; Matthew E Larkum et al., 2009; Sandler et al., 2016). Using the AFL-TMP system to perform highly localized ablation (**Fig. 3**), we can cut an individual dendritic branch and examined the consequences to other sibling branches. In this experiment, we ablated the apical dendrites (10-250 μm) of layer 5 pyramidal neurons in *Thy1*-YFP H line transgenic mice. Time series of fluorescence images were captured at the targeted sites after dendrotomy to evaluate the structural integrity of dendrites around the target sites.

We first used the amplified femtosecond laser to accurately cut a superficial (10-50 μm below the cortical surface) dendritic branch (#1) and found that this cutting did not compromise the morphological structures of its sibling dendritic branch (#2) and a more distant sibling branch (#3) from another primary dendrite (**Supplementary Fig. 6a-c**). Then when the primary dendritic branches deeper (100-250 μm under the pia) in the mouse cortex were severed (**Fig. 5a-d**), no noticeable morphological changes were observed in the apical branches of the targeted primary dendrites or their sibling counterparts at least 1 hour after laser irradiation (**Fig. 5d-f**). The apical branches of targeted primary dendrites did degenerate gradually and disappeared 1-day post laser irradiation (**Fig. 5d**, 1 d).

**Figure 5.**
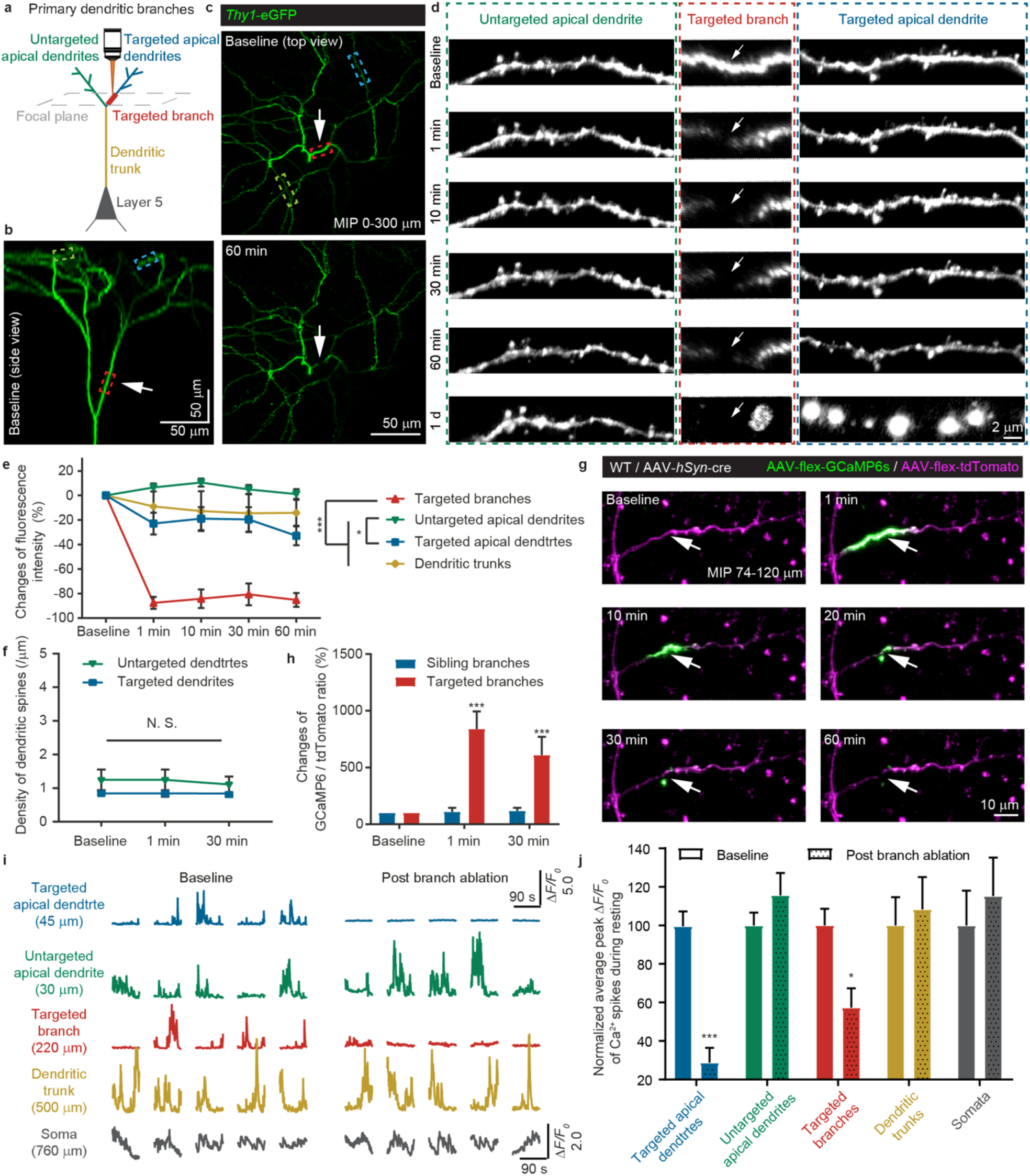
AFL-TPM allows precise dendroctomy at different anatomical locations. a. Schematic of AFL-TPM-mediated ablation of the primary dendritic branch (100 – 250 μm, red) of layer 5 pyramidal neurons. Blue, apical dendrites from the targeted branch; green, untargeted apical dendrites; yellow, dendritic trunk; black, layer 5 soma. Grey dotted box indicates the focal plane for laser irradiation. **b**, The side view of a three-dimensional reconstructed image showing *Thy1*-eGFP labeled apical dendrites of the layer 5 pyramidal neuron. The color-coded dotted frames denote the corresponding three dendritic branches in **a**. Scale bar: 50 μm. **c**, The top view of dendritic branches in **b** before and after (60 min) the primary dendritic truck was severed by the AFL-TPM (0.10 μJ). Images depth: 0-300 μm. Scale bar: 50 μm. **d**, Changes in eGFP signals of the apical dendritic branches in **a-c** over time after the primary dendritic trunk was severed by AFL-TPM. The targeted branch (arrows, red dotted box in **b,c**) showed reduction of eGFP signals immediately after laser irradiation, while the superficial dendrites remained structurally intact for a few hours. Both the targeted branch (middle) and its distal apical dendritic tuft (right) were completely degraded, while the untargeted apical tuft (left) remained intact one day later. Scale bar: 2 μm. **e**, The fluorescence of the targeted branch decreases significantly (*P* < 0.0001) after AFL-TPM-mediated ablation, while other superficial dendrites remained unchanged over a time course of 60 min post laser irradiation. A significant difference is shown between the targeted and untargeted tufts (*P* = 0.011). *n =* 7 target branches, 18 apical dendrites, 5 trunks from 4 mice. Dunnett’s multiple comparisons test. **f**, No significant difference in the density of dendritic spines between the targeted and untargeted apical dendrites. *P* = 0.1. *n* = 114 spines, 5 views from 4 mice. Kolmogorov-Smirnov test. **g**, Representative images showing AFL-TPM irradiation-associated damages were confined within the targeted dendrites. Decreased tdTomado (magenta) and increased calcium (green) signals were observed only in the targeted branch but not in the sibling branch post laser irradiation (0.20 μJ). Scale bar: 10 μm. **h**, An increase in the ratio of GCaMP6 to tdTomato intensity was observed only in the targeted branches (baseline vs. 1 min, *P* < 0.0001; baseline vs. 30 min, *P* = 0.0002). *n =* 5 paired dendrites from 3 mice. Dunnett’s multiple comparisons test. **i**, Examples of calcium transients before and after cutting the primary dendritic branch (at 220 μm) of an individual pyramidal neuron and monitor apical dendrites (at 30 and 45 μm), trunk (at 500 μm), and soma (at 760 μm) during resting. **j**, Summary of average peak Δ*F/F*_*0*_ of the different part of L5 pyramidal neuron before and after cutting primary dendritic branches during resting (Targeted apical dendrites baseline vs. post ablation, *P* < 0.0001; targeted branches, baseline vs. post ablation, *P* = 0.0314). *n =* 25 trials from 5 mice. Sidak’s multiple comparisons test. Data are mean ± s.e.m. **g, f, h, j**, * *P* < 0.05, *** *P* < 0.001.

Next, we used both GCaMP6 and tdTomato viruses to label the apical dendrites of layer 5 pyramidal neurons to further examine whether the AFL-TPM-induced functional damage only occurred in the branches of the targeted dendrites. We observed a burst of GCaMP6 fluorescence in the targeted dendritic branches rapidly (**Fig. 5g**, 1 min) after laser irradiation, which gradually decreased to the undetectable level over 1 hour (**Fig. 5g,h**). No apparent calcium elevation was observed in the sibling dendritic branches (**Fig. 5g,h**). In addition, after cutting the primary branches (target branch in **Fig. 5a**) of layer 5 pyramidal neurons, we found that the spontaneous calcium spikes in targeted branches and their apical dendrites decreased significantly, while the untargeted apical dendrites from other primary branches, trunks, and somata remained relatively unchanged during resting (**Fig. 5i,j**).

Taken together, these findings indicate that the AFL-TPM system can be used effectively to cut one dendritic branch without affecting the integrity of its sibling dendrites. In addition, ablation-induced calcium elevation in individual dendritic branches was maintained within the same branches, suggesting that different dendritic branches of the same neurons undergo changes independently.

## Discussion

In this study, we developed an amplified femtosecond laser-mediated *in vivo* cell ablation technology. Utilizing an iterative convergence control, we were able to ablate individual cells precisely and effectively at the subcellular level in the living mouse cortex, without altering the structural and functional integrity of nearby tissues. By targeting SST-expressing interneurons in layer 2/3 of mouse neocortex, we demonstrated that ablating few SST-expressing interneurons has a significant inhibitory effect on the activity of nearby neurons. Furthermore, cutting a dendritic branch has no obvious effects on the integrity of its sibling branches, suggesting dendritic branches are independent structural and functional units.

Currently, Ti:sapphire laser is a widely used tool for the ablation of selective neuronal cells(Hayes et al., 2014; Orger et al., 2008; Pozner et al., 2015). However, it requires prolonged scanning time to ablate cells due to the low energy in each pulse. The cumulative thermal effect caused by continuous illumination might lead to a secondary damage to the surrounding tissue of the targeted cells(Bauer et al., 2015; Di Niso et al., 2014; Jasiński, 2018). Furthermore, the two-photon Ti:sapphire laser has limited applications in deep cortical tissues. In contrast, the amplified femtosecond laser can generate a single pulse with enough power (20,000 times higher than that of Ti:sapphire laser) to ablate targeted cells without the need of continuous illumination(Krüger and Kautek, 2012; Strickland and Mourou, 1985). Previous studies have used such amplified femtosecond laser to cut the axons of worms(Bourgeois and Ben-Yakar, 2008; Chung and Mazur, 2009; Gabel et al., 2008; Guo et al., 2008; Morris et al., 2013; Wu et al., 2007; Yanik et al., 2004), ablate cells in cultures(Hamad, 2016)(Watanabe et al., 2004)and brain slices(Tsai et al., 2009), and model stroke by occluding cerebral vessels *in vivo*(Blinder et al., 2012; Nishimura et al., 2007, 2006; Tsai et al., 2003). However, it has not been used for ablating individual cells in the brain in a spatia-temporal controlled manner. By developing an iterative convergence power and Galvo control system, we were able to control precisely both the power of each amplified femtosecond laser pulse and the targeted ablation position in a 3D volume. We demonstrated, for the first time, that amplified femtosecond laser is capable of targeting and ablating cells in the cortex of awake behaving mice with minimum collateral damage to nearby tissues (within 1 μm distance).

The success of the AFL-TPM system depends on the high peak power of each pulse to induce necessary nonlinear absorption, while maintaining adequately low average powers to avoid linear heating(Tsai et al., 2009). In addition, the shorter pulse duration can lead to larger peak power under a constant pulse energy(Hamad, 2016). Because the pulse of amplified femtosecond laser we used (35 fs, **Supplementary Fig. 1b**) was much shorter than the previous studies(Blinder et al., 2012; Bourgeois and Ben-Yakar, 2008; Chung and Mazur, 2009; Gabel et al., 2008; Guo et al., 2008; Morris et al., 2013; Nishimura et al., 2007, 2006; Tsai et al., 2009, 2003; Wu et al., 2007; Yanik et al., 2004) (around 100 fs), this enables us to ablate individual cortical cells deep *in vivo* with one laser pulse.

The amplified femtosecond-based cell ablation also offers several advantages over other ablation methods. First, although chemical-, genetic-, optogenetic-based ablation strategies can be used to target cells embedded deep in tissues(Cheng et al., 2016; Damisah et al., 2017; Makhijani et al., 2017; Tertilt et al., 2005), a relatively long period of time (hours to days) is required to ensure the death of targeted cells through apoptosis before any changes could be monitored. Because neural network may compensate for the loss of functions of ablated cells, the changes of neural networks in response to apoptosis-based targeted cell ablation could be masked(Barron et al., 2017; Driscoll et al., 2017; Dunn, 2015b). In contrast, since the amplified femtosecond laser could deactivate targeted cells instantaneously, it would allow us to detect immediate changes in neural network without the complication of compensatory responses post ablation. Second, genetic and optogenetic-based ablation methods could be limited by the lack of specific promoters for cell type-specific gene expression. On the other hand, the AFL-TPM system allows ablation of cells either expressing fluorescent proteins or labeled by fluorescent dyes, which does not require specific promoters. Third, apoptosis-based ablation target entire cells, while AFL-TPM could ablate part of an individual cell, allowing a better understanding of dendritic or axonal functions(Cichon and Gan, 2015). By performing precise dendrotomy of apical dendrites using the AFL-TPM, we observed that the increased calcium was confined within the injured dendrites and did not spread to their neighboring branches. These findings, in line with those *in vitro* studies(M. E. Larkum et al., 2009), suggest each dendritic branch as an independent functional unit.

It is important to mention that the optogenetic manipulation coupled with a spatial light modulator (SLM) has recently been applied to neurons infected with red-shift light-sensitive ion channels. This method could either activate or inhibit neuronal population at the single cell resolution, thereby modulating the excitability of neural networks(Mardinly et al., 2018). However, manipulating neuronal activity with the SLM based optogenetic method relies on the level of light-sensitive opsin expression. The amplified femtosecond laser ablation method offers a simple and alternative means to investigate the function of cells without the need to expressing light-sensitive opsins in specific cell types. Consistent with previous studies using wild field optogenetic methods to inhibit SST-expressing interneurons(Fu et al., 2014; Gentet et al., 2012) (Adler et al., 2019), we deactivated individual SST interneurons using the amplified femtosecond laser ablation method, and found that ablation of a few SST cells in the primary motor cortex increased activity of surrounding neurons during treadmill running. Thus, the AFL-TPM ablation method can serve as a simple and effective means to examine the functions of specific neuronal types *in vivo*.

In addition to targeting individual neurons and neuronal populations in the living brain, the amplified femtosecond laser-based platform we developed here is also capable of targeting non-neuronal cells such as astrocytes and microglia (**Supplementary Fig. 7**), making this tool more flexible for studying functions of various cell types *in vivo*. In combination of transgenic lines to labeled specific cell types, we envision this method will greatly facilitate studies of cell functions in the complicated neural network in the future.

## Materials and Methods

### Mice

The wild-type (WT) C57BL/6 mice and transgenic mice *Thy1*-YFP in layer 5 pyramidal neurons (H line), *Thy1*-eGFP in pyramidal neurons (M line), *Cx3cr1*-GFP mice expressing eGFP in microglia cell (*Cx3cr1*^*tm1Litt*^*/J*), and SST-IRES-Cre mice (*Sst* ^*tm2*.*1(cre) Zjh*^*/J*) were purchased from the Jackson Laboratories. Mice were housed in Bindley Bioscience Center Animal Facility at Purdue University. All the one- or two-months old animal procedures were performed under the institutional guidelines.

### Virus injection

To prevent brain edema, dexamethasone sodium phosphate (BP108, 1.32 mg/kg, Sigma-Aldrich) were intraperitoneal (i.p.) injected 30 minutes before virus injection. The mice were anesthetized with 3% isoflurane and maintained with 1.5% isoflurane (Matrx VIP 3000 isoflurane vaporizer, Midmark) until the end of surgery. The head of the mouse was fixed to the stereo locator (Model 940, David KOPF Instruments) after the toe unresponsiveness check. Before viral injection, glass electrodes with tip size less than 0.5-micron were prepared by glass micropipette puller (P-97, Sutter Instrument), back-filled with mineral oil (16242, Cagrille Labs), and front-loaded with a solution containing AAV viruses. 200-300 nl solution containing Adeno-Associated Viruses (AAV) was injected in the cortex of each mouse with a Nano injection pump (Nanoject II, Drummond Scientific Company) with a speed of 30 nl/min. The viruses were injected into the primary motor cortex (0.5 mm anterior and 1.2 mm lateral from bregma, depth of 200-300 μm for layer 2/3, 500-600 μm for layer 5)(Cichon and Gan, 2015). After injection, the glass electrode stayed in the brain for 5 min to ensure that the spread of injected viruses. The skin was stitched, and mice were placed back to their home cages after they fully recovered from anesthesia. Buprenorphine was applied at 0.1 mg/kg (i.p.) for three days after surgery avoiding pain relief.

For morphological assessment, *Cx3cr1*-GFP, *Thy1*-YFP-H, and *Thy1*-eGFP-M mice were used for imaging. For functional studies, GCaMP6 was expressed with Adeno-Associated Virus (AAV) that was injected into the primary motor cortex of mice. The injected viruses were ordered from Addgene included: AAV1-*Syn*-GCaMP6s-WPRE-SV40 (100843-AAV1, 1×10^13^), AAV1-*hSyn*-Cre-WPRE-hGH (105553-AAV1, 1×10^13^), AAV5-*gfaABC1D*-lck-GCaMP6f (52924-AAV5, 7×10^12^), AAV9-*pCAG*-FLEX-tdTomato-WPRE (51503-AAV9, 1×10^13^), AAV1-*CAG*-Flex-GCaMP6s-WPRE-SV40 (100842-AAV1, 1×10^13^). GCaMP6s driven by the *Syn* promoter were injected for experiments described in **Fig. 2e-g** and **Supplementary Fig. 2**. For functional studies of an individual nerve cell or the entire neural network, the cre-dependent flex-tdTomato and *Syn-*driven GCaMP6s viruses were co-injected to the primary motor cortex of SST-cre mice in **Fig. 4** and **Supplementary Fig. 5**. To simultaneous imaging of calcium and neuronal structure, two cre-dependent DIO viruses (tdTomato and GCaMP6s) were used in **Fig. 5g-j**. In order to reduce the background fluorescence, the flex viruses were mixed with highly diluted (1: 25,000) cre virus. The fully mixed viruses were co-injected into layer 5 for labeling pyramidal neurons. The cytoplasm calcium of astrocytes was detected by *gfaABC1D* derived GCaMP6f in **Supplementary Fig. 7**.

### Surgical procedures

Before surgery, mice were anesthetized with 3% isoflurane and maintained with 1.5% isoflurane until the end of the operation. After the mice showed no response to a toe pinch, povidone-iodine and 70% alcohol were applied to disinfect the head skin. Ophthalmic ointment was applied to both eyes to prevent desiccation. The mice were transferred to a 37°C heating pad (TCAT-2LV, Physitemp Instruments) to maintain their body temperature. All surgeries were performed under an industrial microscope (SZX7, 1.25 × achromatic objective lens, Olympus) with fiber optic cold light sources (V-Lux 1000, Harvard Apparatus). To reduce the movement of brain tissue during imaging, dental cement was used to secure two custom-made head bars to the skull after removing the skin. A high-speed cranial drill (K107018, Foredom Electric Co.) was used to polish the skull above the mouse motor cortex based on stereotaxic coordinates. When the skull became thin enough to visualize the target brain area, microscopic tweezers were used to carefully lift the skull, while avoiding damages to the dura and blood vessels. The artificial cerebrospinal fluid (ACSF, 119 mM NaCl, 2.5 mM KCl, 1.3 mM MgCl_2_, 26 NaHCO_3_, 1 mM NaH_2_PO_4_, 2.5 mM CaCl_2_, 10 mM Glucose, and pH adjusted to 7.2) was applied to the exposed cerebral cortex and a 0# glass with a diameter of 3 mm (CS-3R-0, Warmer Instruments) was used to cover the exposed cortex and tightly glued to the skull. Upon completion of the surgery, mice were either returned to the cage or imaged immediately.

### Labeling procedures

Fluorescent dyes were applied immediately after the skull was opened in some experiments. Propidium Iodide (PI, 0.5 mg/ml(Antkowiak et al., 2013; Wilde et al., 1994), Anaspec) and Hoechst (33342, 0.04 mg/ml(Damisah et al., 2017; Latt et al., 1975), Thermos fisher) were applied together for 10 mins to label the nuclei of dead and live cells, respectively. Sulforhodamine 101 (SR101, 200 mM(Nimmerjahn et al., 2004), Sigma-Aldrich) dissolved in ACSF was briefly (3-5 minutes) applied on the exposed topical surface to label cortical astrocytes. The cortex was then rinsed with ACSF for 2-3 times. After the dyes were applied, the exposed cortex was covered by a 3-mm 0# glass (CS-3R-0, Warmer Instruments) that was tightly glued to the skull. For the vascular labeling experiments, Dextran & Texas Red (70,000 MW, 5%, 50 mg/kg(Nimmerjahn et al., 2004), Thermo Fisher) was used by orbital injection.

### Two-photon imaging system

We built a two-photon microscope (TPM) for structural and functional imaging before and after cell ablation (**Fig. 1a**). The imaging beam was from a 140 fs 80 MHz repetition rate Ti:sapphire oscillator (Chameleon Ultra II, Coherent). The overall imaging beam dispersion was compensated by an automated prism-based pulse compressor (Chameleon PreComp, Coherent). The imaging beam intensity was controlled by an electro-optic modulator (M350-80, Conoptics). We expanded and collimated the ablation beam and the imaging beam, and combined the beams with a dielectric beam splitter with low group delay dispersion (OA037, Femtooptics). The combined beams entered a 5 mm diameter *xy* Galvo mirror pair (6215H, Cambridge Technology). We used two customized telecentric relay scan lenses (*f* = 125 mm and *f* = 500 mm) to image the Galvo onto the back focal plane of a high NA low magnification water dipping objective (Nikon 25x NA 1.1). The two-photon excited fluorescence was imaged by customized high etendue optics onto the aperture of GaAsP PMTs (H10770PB-40, Hamamatsu). We used three color channels for cyan, green and red emissions (ET460/36m, Chroma; FF01-520/70-25, Semrock; FF01-607/70-25, Semrock). A high-precision and high-repeatability stage (M-VP-25XA-XYZR, Newport) was utilized to control the animal position. For volumetric imaging, we used the Galvo for transverse scanning and used the stage for axial movement. The two-photon imaging was controlled by *Scanimage* (Vidrio Technologies). The two-photon imaging laser was tuned to 775 nm for Hoechst 33342 imaging, and 930 nm for Propidium Iodide, eGFP, YFP, GCaMP6, tdTomato, Sulforhodamine 101 and Dextran & Texas Red imaging.

### Laser-mediated ablation with iterative convergence position control

The ablation laser source was a 3 W, 35 fs (**Supplementary Fig. 1b**), 800 nm, 10 kHz repetition rate regenerative chirped pulse amplifier (Spitfire Pro, Spectra Physics). With its high pulse energy, we could afford to use 50% ultrafast beam splitter (OA037, Femto optics) to combine it with the imaging beam. As our goal is to achieve high-precision subcellular ablation, we need to ensure long-term alignment stability. Experimentally, we utilized one of the two outputs from the beam splitter for biology measurement and used the second output for position sensing. Specifically, we focused the combined beam with a lens onto a camera (DFK 72BUC02, Imaging Source). Thus, from the relative focal spots distance (a 2D vector), we could accurately determine the relative angular drift between the imaging beam and the ablation beam, which was automatically measured before each ablation to accurately account for the beam drift.

To accurately control the Galvo to deliver a single pulse to the target location, we developed an iterative convergence control method. Through the experimental test, we found that the built-in feedback signal of the slow Galvo scan axis could serve as a reliable position feedback signal. The fast axis feedback had a speed-dependent delay, which was unreliable (**Supplementary Fig. 1c**). To utilize only the slow axis signal, we let the *xy* Galvo scanners took turns to serve as fast and slow axis and combined the image results to extract the position data for the two scanners. Through calibration, we could determine the linear relationship between the Galvo driving voltage and the Galvo feedback, which however had a very slow variation over time. To eliminate such error, we iteratively drove the Galvo and measured the feedback. After each iteration, we analyzed the difference between the desired and the measured Galvo feedback position and subsequently revised the Galvo driving signal. Typically, we used three iterations to convert the desired feedback signal to the actual driving voltage. All these technical procedures (drift compensation, slow Galvo feedback, and the iterative convergence-based control) lead to the 3D high-precision ablation demonstrated in this work.

To provide accurate timing control for single-pulse ablation, we utilized an electro-optic modulator (M350-80, Conoptics) with high-extinction-ratio to gate single pulses out of the 10 kHz pulse train (**Supplementary Fig. 1a**). The modulator’s driving signal was triggered by the regenerative amplifier’s 10 kHz clock. To control the pulse energy over a wide dynamic range, we utilized achromatic half-wave plate (WP, 10RP52-2B, Newport) and polarizer (GL15-B, Thorlabs) as the secondary power control. The ablation pulse energy was recorded in real-time by an amplified photodiode (PDA36A, Thorlabs).

For calibration, we dried a layer of 1 μm yellow-green fluorescence microspheres (F8852, Thermo Fisher) on a cover glass. The axial focus positions of the imaging beam and ablation beam were measured by using each beam to image the same bead layer. For the *in vivo* AFL-TPM ablation experiments, we linearly increased the pulse energy and measured the average fluorescence intensity of the target center immediately. When the average structure fluorescence intensity were reduced to 10% or less or when the calcium signals was increased to sufficiently light up the entire soma, we stopped applying further energy increase. For the comparison experiment using the 80 MHz 140 fs Ti:sapphire imaging beam for ablation, we raster scanned a small area (1∼5 μm^2^) for 2-5 sec. Typically, we need to use the maximum imaging beam power to generate thermal ablation(Buffelli et al., 2007; Fu et al., 2014; Hayes et al., 2012; Li et al., 2018; Pozner et al., 2015; Rompolas et al., 2012).

### Treadmill training

A custom free-suspended treadmill (9 × 15 cm) was used for motor training. The treadmill allowed head-fixed mice to move their forelimbs freely on the treadmill track during the forward running task(Cichon and Gan, 2015; Li et al., 2017). To minimize the movement of images during the imaging process, any moving parts (*i*.*e*., motors, belts, and drive shafts) of the treadmill was not in contact with the electric stage or microscope. During imaging, the head of the experimental mouse was placed on the electric stage (M-VP-25XA, Newport) and fixed with screws. Both head-fixed mice and the stage were not in contact with the two-photon microscope. During the running training, the treadmill track was driven by a motor (Model 2l010, Dayton) under the 10 V DC power condition. Each running trial lasted for 60 seconds, with at least 90 seconds or longer break interval before the next trial began (**Supplementary Fig. 2a**).

### *In vivo* two-photon imaging and data analysis

For morphological imaging experiments, volumes were collected from the primary motor cortex region of each mouse, with a field of views of 235 μm (Zoom 3) or 142 μm (Zoom 5). The two-photon imaging frame rate was 0.36 fps with an image size of 1024 × 1024. The interval between each slice in the Z direction was set at 1, 2, 5 μm, depending on the specific experiments. The process of side view was reconstructed by the 3D stack function of *Image J* software, and all Z overlay images were projected with the maximum fluorescence for each slice (maximum intensity projection, MIP).

For calcium imaging experiments, data were collected from layer 2/3 of the primary motor cortex. The two-photon imaging frame rate was 0.72 fps with an image size of 1024 × 1024. Under resting conditions, the treadmill was turned off and two-photon imaging was conducted for 90 s. The running trial contained three periods: started with a 12-s resting time, followed by 60-s running time, and ended with an 18-s post-resting time. Both the resting and running periods were repeated 3-5 times.

All the time-lapse images with the same views were registered with the averaged image using two-dimensional (2D) cross-correlation for motion correction. For double-labeling experiments, motion compensation was obtained by using 2D cross-correlation with the morphological dataset and then applied to GCaMP6 dataset. For GCaMP6 only dataset, the average of the first trial was used to correct the whole GCaMP6 dataset. Changes of fluorescence (Δ*F/F*_*0*_) were quantified from regions of interests (ROI), corresponding to visually identifiable somata. The minimum 10% of each cell’s fluorescence was defined as *F*_*0*_ for the resting quantification(Mardinly et al., 2018), and the average fluorescence within a 3 s window before running was selected as *F*_*0*_ for the running quantification(Cichon and Gan, 2015). The threshold for detecting calcium spikes was > 0.5 Δ*F/F*_*0*_. The full width at half maximum (FWHM) of each calcium spike was measured as the calcium spike duration(Li et al., 2017). The frequency was defined as the number of calcium spikes in every second; the peak amplitude was defined as the maximum value of each calcium peak. The total calcium activity was defined as the accumulated area under the curve per second. All the above data for calcium spikes did not overlap with other spikes.

### Statistics

All the data in this study are represented as mean ± standard error. All the data were performed with the Shapiro-Wilk normality test. For the data passed normality test (*P* > 0.05), the two-way ANOVA Dunnett’s multiple comparisons test and Sidak’s multiple comparisons test were selected for comparing multiple groups. During the analysis process, pairing or non-pairing detection was selected according to the specific circumstances of each experimental condition and was clearly indicated in each figure legend. For experimental data that did not conform to a normal distribution, the non-parametric Mann Whitney test or Kolmogorov-Smirnov test was performed. *P* < 0.05 is recognized as statistically significant. All statistical analyses were performed using *GraphPad Prism*. No results of the successful acquisition from images and measurements were excluded and filtered. The experiment did not include randomized and blinding experiments. At least three mice were used in each experiment. *P* values, *n*, and the statistical tests were described in the figure legends and supplementary table.

## Data and codes availability

All the data and codes described in this study are available publicly.

## Acknowledgments

We thank all the members in the Gan Laboratory and the Cui Laboratory for their constructive discussion on this project. This work supported by NIH grant U01NS094341, U01NS107689, and the research fund provided by Purdue University to the Cui lab. Z.-Y.C. acknowledge the support from the Ph.D. abroad visiting scholar program of the Nanchang University. M.C. acknowledge the scientific equipment support from HHMI.

## Author contributions

W.-B.G. and M.C. initiated the project. M.C. designed and implemented the overall imaging and ablation system. Y.-Y.H. performed the system calibration, wrote the ablation control program and data processing code. B.-W.W. maintained, optimized and calibrated the ablation laser source. Z.-Y.C. and W.-B.G. designed the experiments. Z.-Y.C. and Y.-Y.H. performed the experiments and analyzed the data. Z.-Y.C., Y.-Y.H., M. C, and W.-B.G. wrote the manuscript.

## Competing Financial Interests

The authors declare no competing financial interests.

## Supplementary Information

**Supplementary Figure 1.**
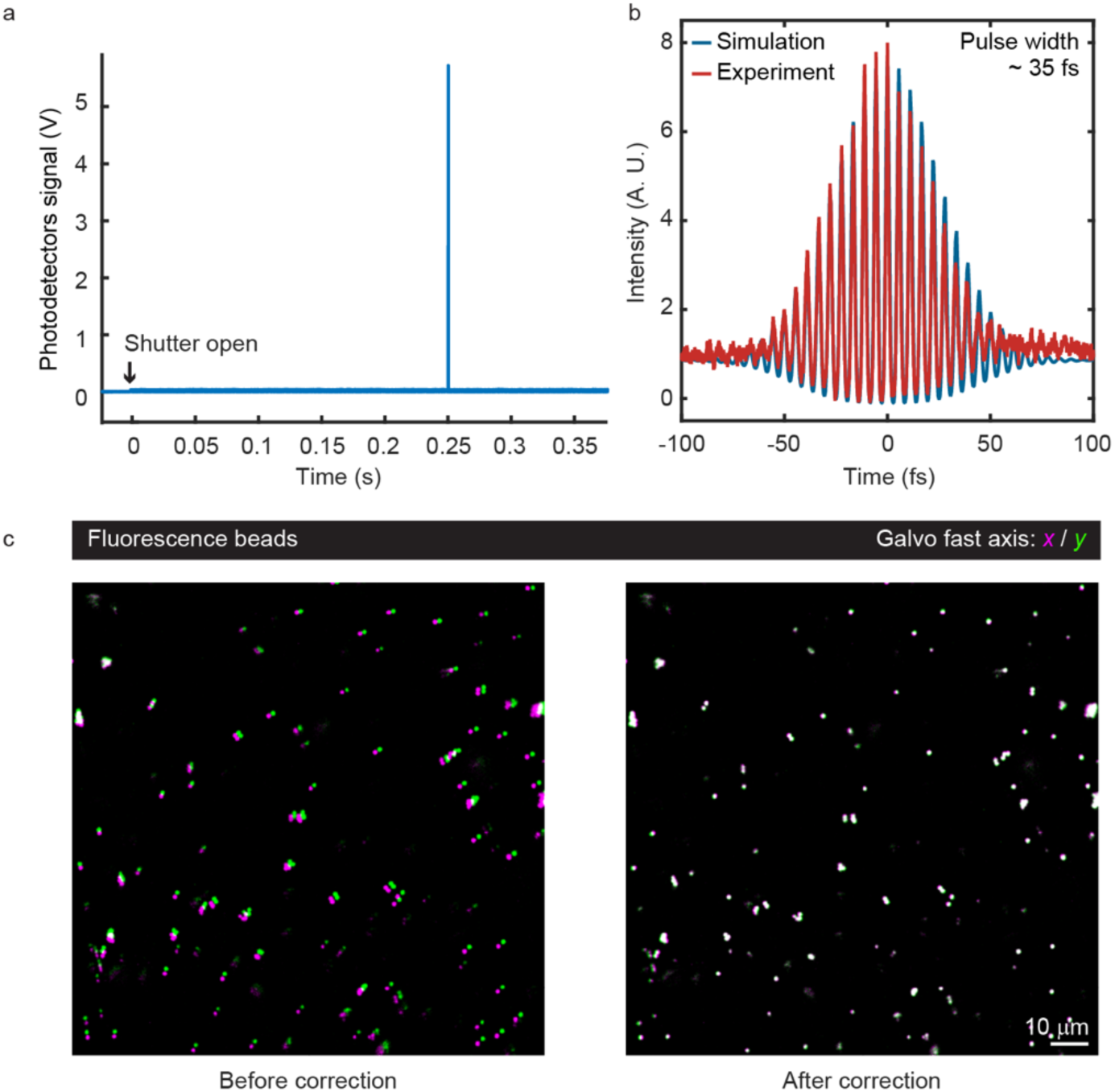
Pulse parameters of amplified femtosecond laser and Galvo feedback correction. **a**, During the process of ablation, the single pulse was controlled by electro-optic modulators (EOM in **Fig. 1a**). **b**, The pulse width (35 fs) of amplified femtosecond laser was measured by optical autocorrelation. **c**, The distortion between fast *x* and *y* scanner of Galvo (left) was corrected (right, 1 μm fluorescence beads in 1.5% ager).

**Supplementary Figure 2.**
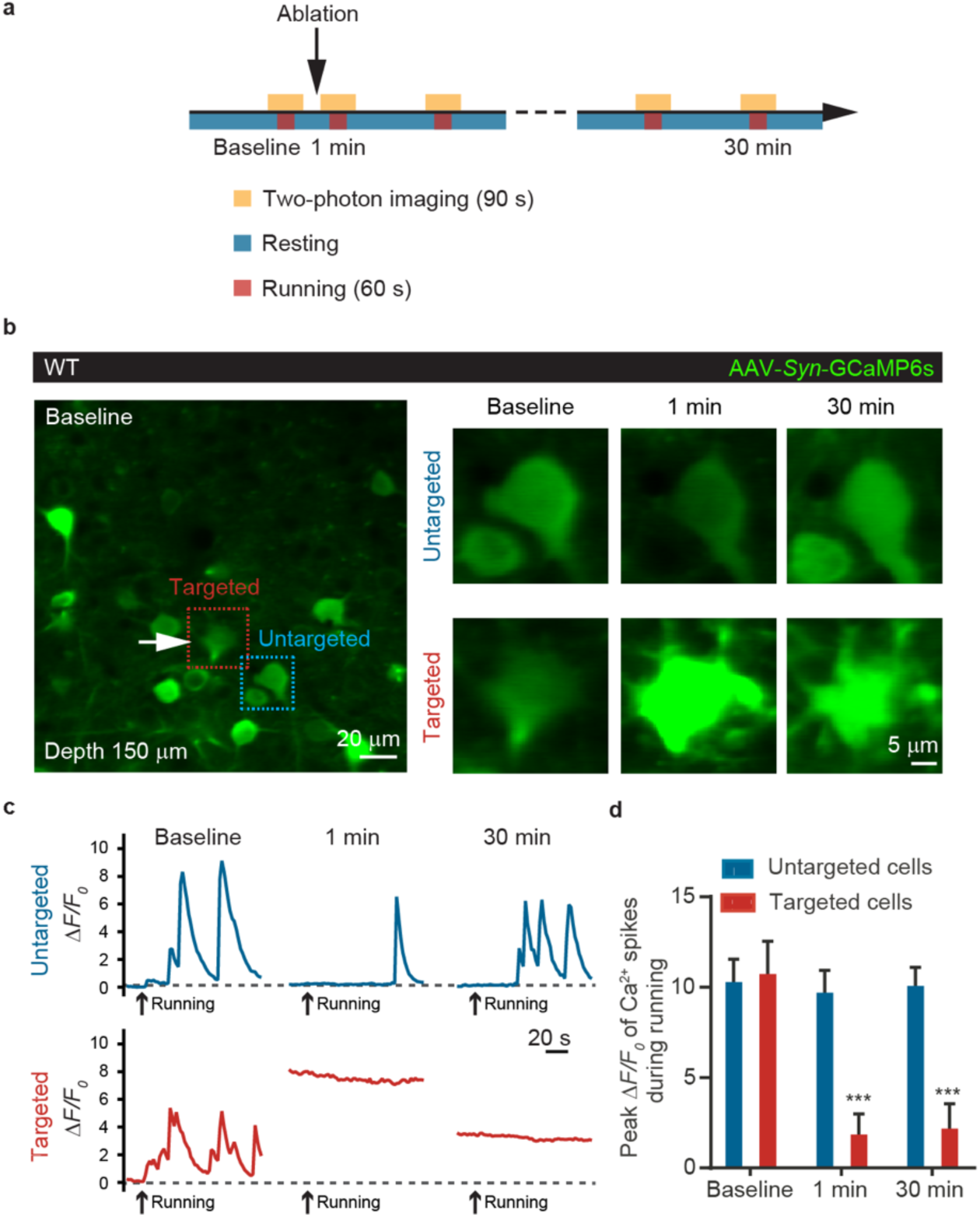
AFL-TPM completely abrogates activity of targeted neurons in the motor cortex. **a**, Schematic showing the time intervals when mice were resting or running and the time points for calcium imaging in the mouse motor cortex. **b**, Two-photon imaging of somatic calcium in layer 2/3 neurons expressing GCaMP6. Calcium signals in the targeted cell (arrow) increased rapidly (1 min) after AFL-TPM irradiation (0.24 μJ) and gradually reduced (30 min). Images depth: 150 μm, scale bars: 20 and 5 μm. **c**, Running-induced calcium changes in the targeted and untargeted neurons before and after AFL-TPM system-mediated ablation. *F*_*0*_ of baseline was used during the Δ*F/F*_*0*_ quantification of 1 min and 30 min. The black arrow denotes the start point of running. Targeted neuron (arrow) showing no response to running-induced calcium activity after AFL-TPM-mediated ablation. **d**, Quantification of the peak Δ*F/F*_*0*_ of GCaMP6 signals during the running state. The peak calcium transients were significantly decreased in targeted cells (1 min: *P* < 0.0001, 30 min: *P* < 0.0001) but not in untargeted cells. *F*_*0*_ was chosen from the averaged 3 s before running of each trial. *n =* 8 pairs of cells from 4 mice. Dunnett’s multiple comparisons test.

**Supplementary Figure 3.**
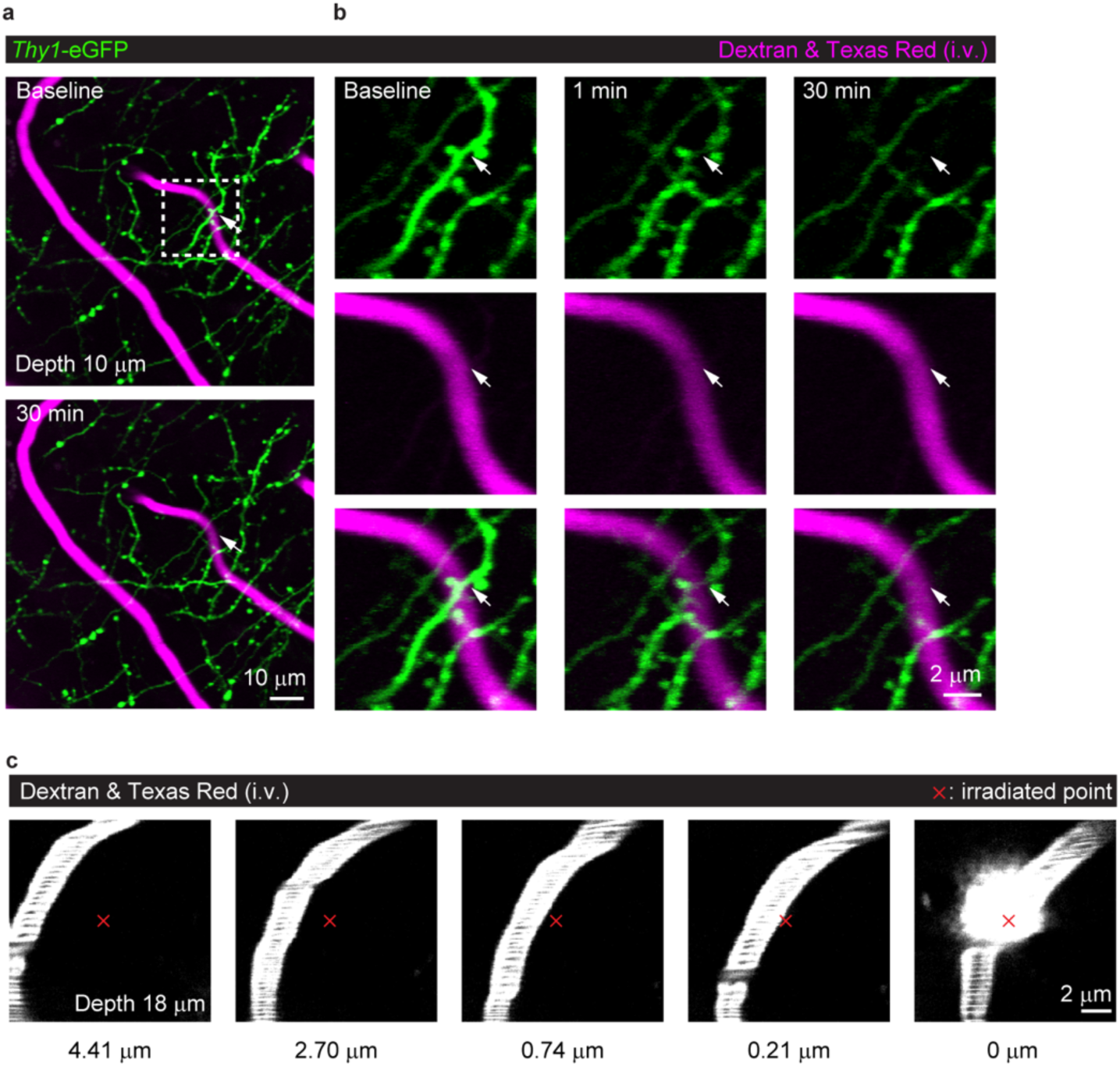
AFL-TPM ablation of dendrites around capillaries didn’t affect the morphology of blood vessels. **a, b**, *In vivo* two-photon imaging of *Thy1*-eGFP labeled apical dendrites (green) and Dextran & Texas red-labeled blood vessels (magenta, i.v.) before and after AFL-TPM ablation. Targeted dendrite (arrow) disappeared after ablation (0.13 μJ), leaving capillaries at the same focal plane intact. Time-lapse images of dendrites and capillaries in the white dotted box (**a**) are shown at higher magnification in **b**. Images depth: 10 μm, scale bars: 10 μm (**a**) and 2 μm (**b**). **c**, AFL-TPM induced capillary breakage only when the irradiated spot was in the center of the blood vessels. Red cross symbols denote the irradiated points (0.09 μJ). Images depth: 18 μm. Scale bar: 2 μm.

**Supplementary Figure 4.**
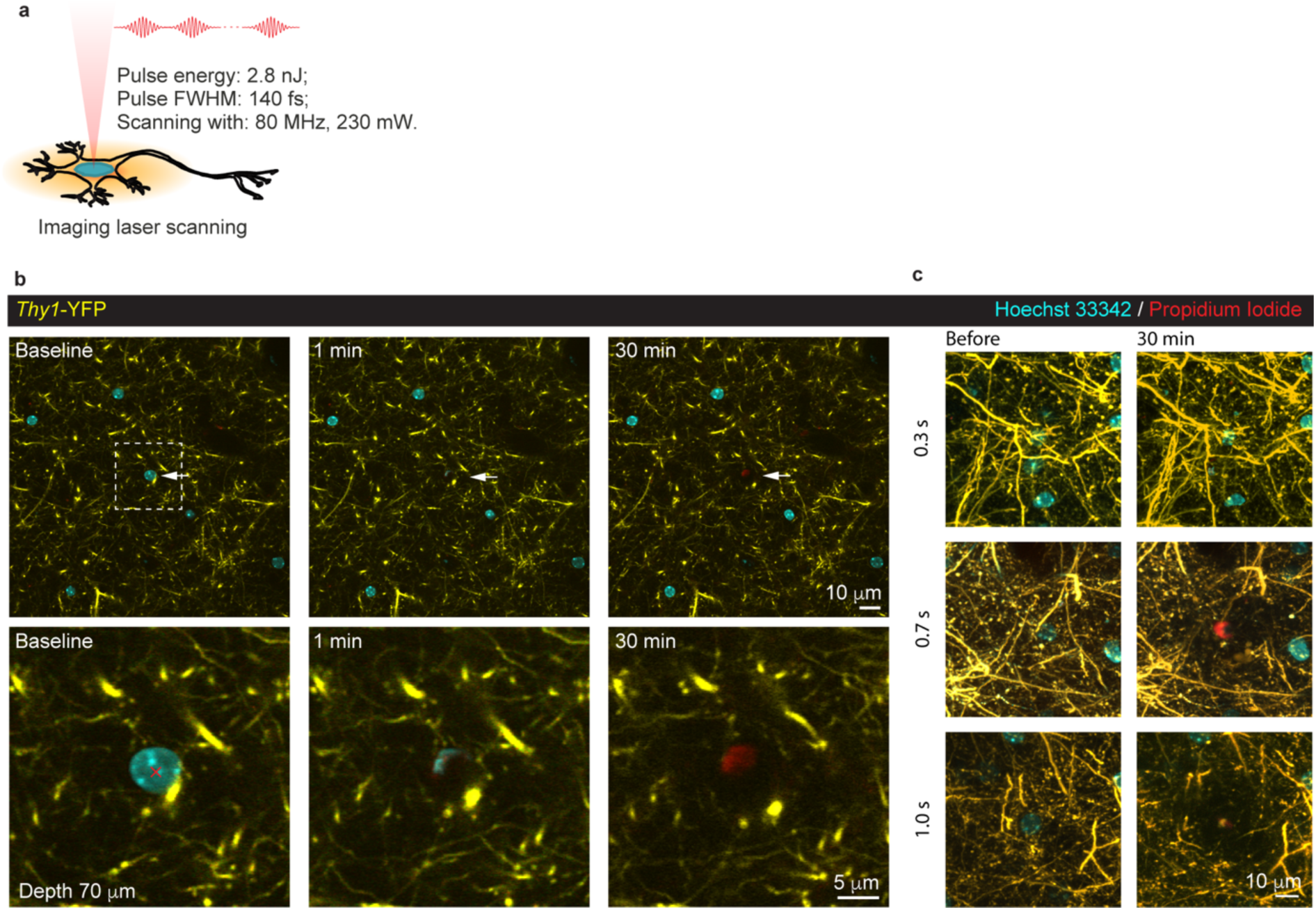
Parking the imaging laser in single point damaged surrounding neuronal structures. **a**, Schematic of two-photon laser parking-mediated cell ablation with multiple pulses. The two-photon laser parked in the nucleus region of interest. The energy for each pulse was 2.8 nJ. **b**, The low pulse power imaging laser parking in single point (red cross, center of targeted nucleus) damaged neuronal structures surrounding the targeted cell due to heat accumulation (110 mW, 5 s). Images depth: 70 μm, scale bars: 10 μm (up) and 5 μm (down). **c**, Comparing of different parking time for imaging laser damage. Scale bars: 10 μm.

**Supplementary Figure 5.**
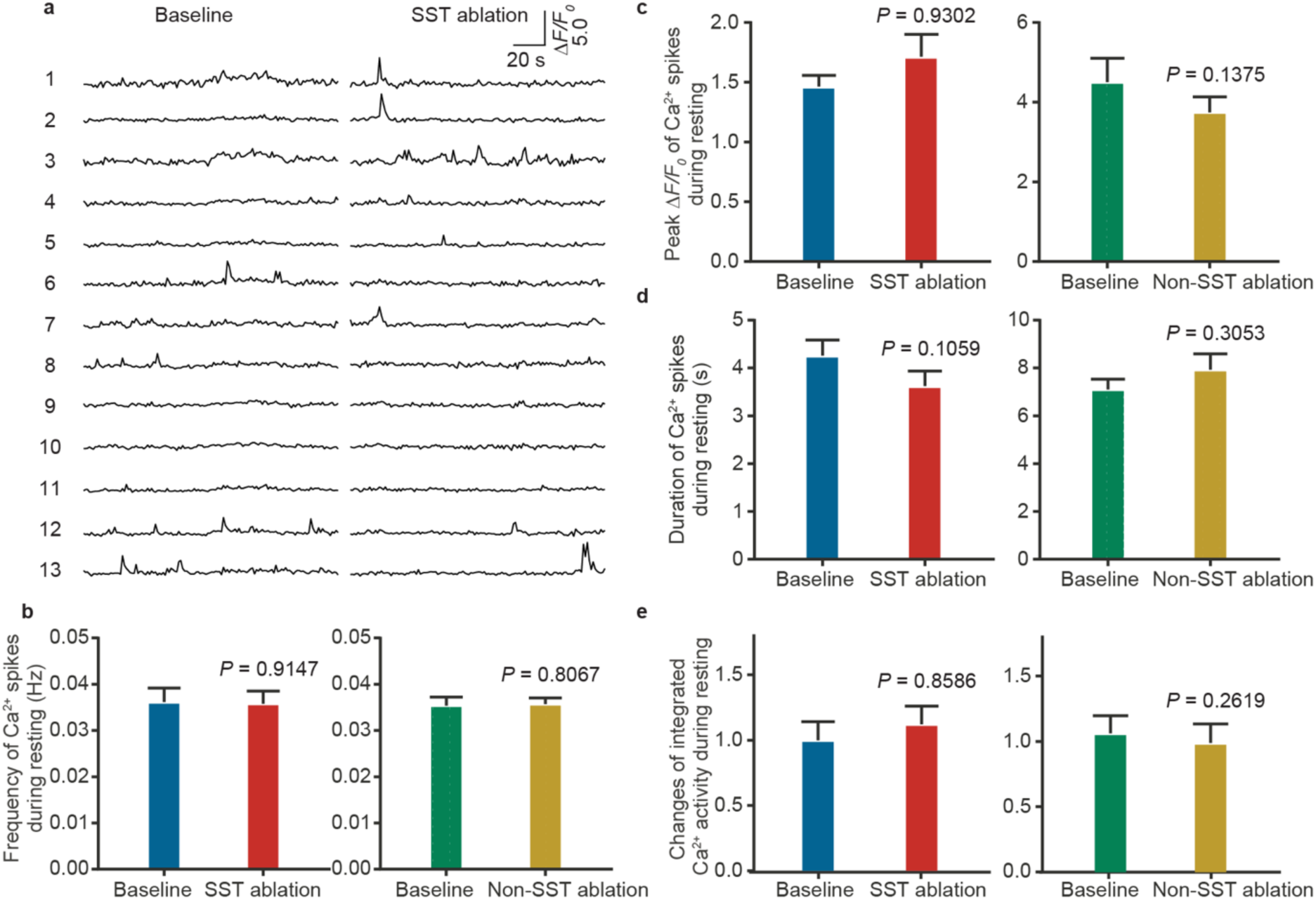
Ablation of SST interneurons or non-SST cell does not affect activity of nearby neurons under resting condition. **a**, Calcium traces of nearby neurons under resting conditions before and after the ablation of SST cells. **b-e**, Frequency, peak Δ*F/F*_*0*_, duration, and integrated calcium activity showing no changes in nearby neurons before and after the ablation of SST cells or non-SST cells. *n =* 49 cells (baseline), and 63 cells (ablation) from 6 mice of the SST group and 137 cells (baseline), and 119 cells (ablation) from 4 mice of the non-SST group, Mann Whitney test.

**Supplementary Figure 6.**
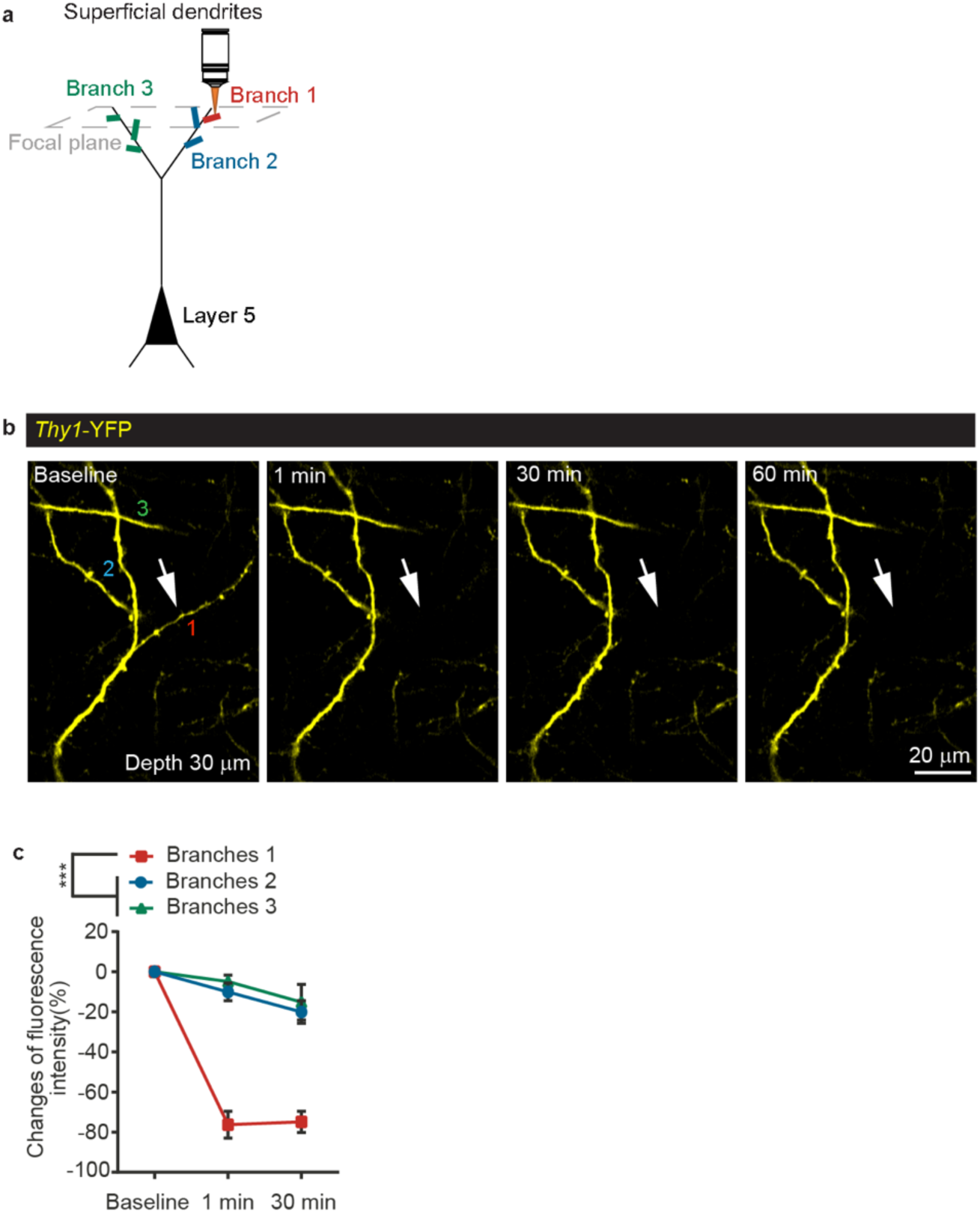
Ablation of an apical tuft dendritic branch did not compromise its sibling branches. **a**, Schematic of AFL-TPM-mediated ablation of apical tuft branches (10-50 μm). Branch 1 (red), the targeted dendrite; branch 2 (blue), the untargeted sibling of branch 1; branch 3 (green), the untargeted dendritic branch from a nearby primary dendritic branch. Grey dotted box denotes the focal plane for laser irradiation. **b**, Representative images showing that AFL-TPM ablated the apical tuft dendrite (arrows) without damaging their sibling dendritic branches (0.05 μJ). Numbers 1-3 denote the three branches defined in **a**. Images depth: 30 μm. Scale bar: 20 μm. **c**, Fluorescence intensity changes in three apical tuft dendritic branches in **b** at different time points post laser irradiation. Branches 1 vs. branches 2 or 3, *P* < 0.0001. *n =* 7, 15, and 9 dendrites from 4 mice. Dunnett’s multiple comparisons test.

**Supplementary Figure 7.**
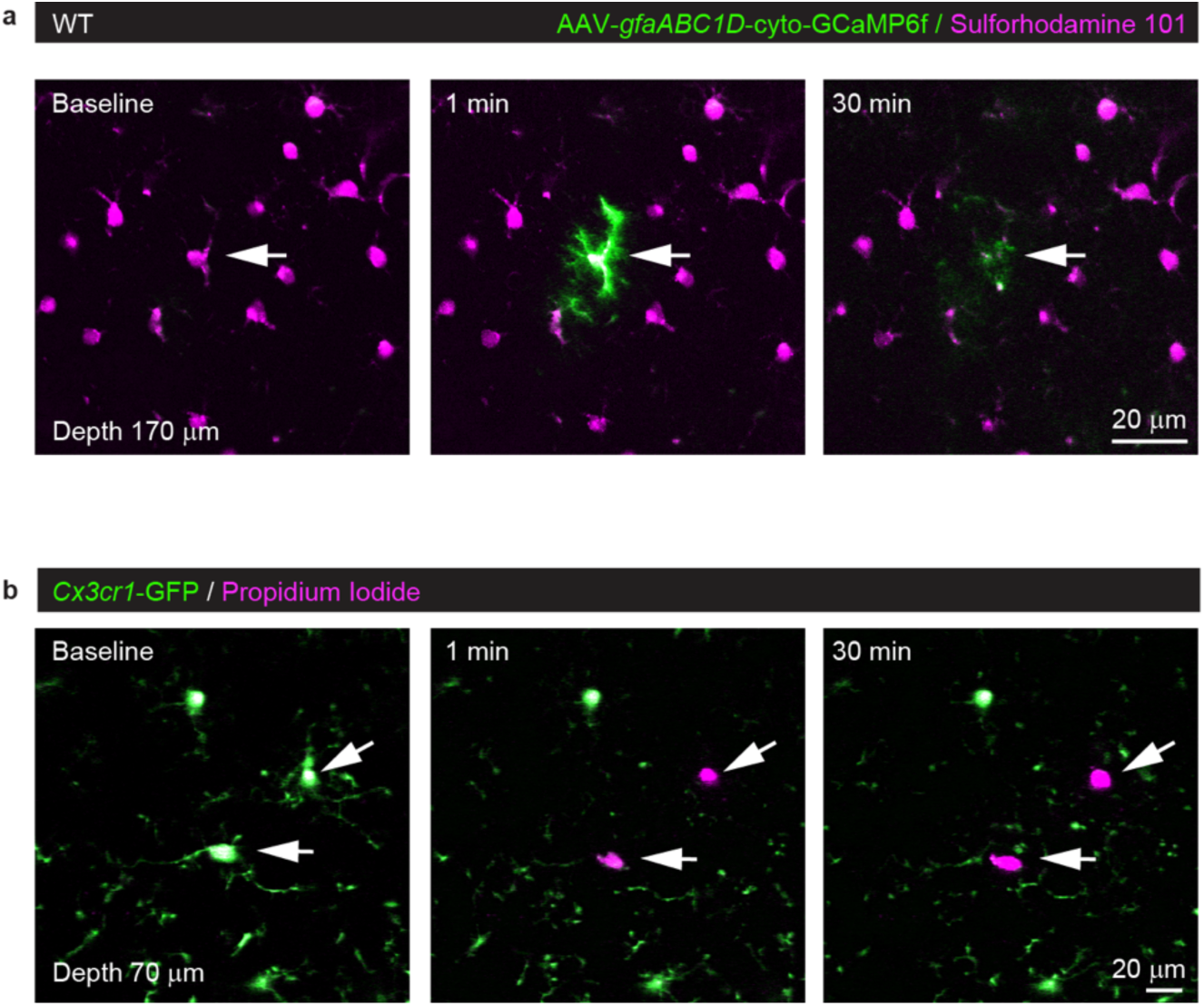
AFL-TPM-mediated ablation of glia cells. **a**, Ablation of Sulforhodamine 101 labeled astrocyte (arrows). **b**, Ablation of *Cx3cr1*-GFP positive microglia with Propidium Iodide (arrows).

## References

Adesnik H, Bruns W, Taniguchi H, Huang ZJ, Scanziani M. 2012. A neural circuit for spatial summation in visual cortex. Nature 490:226–230. doi: 10.1038/nature11526

Adler A, Zhao R, Shin ME, Yasuda R, Gan W-B. 2019. Somatostatin-Expressing Interneurons Enable and Maintain Learning-Dependent Sequential Activation of Pyramidal Neurons. Neuron 102:202-216.e7. doi: 10.1016/j.neuron.2019.01.036

Allegra Mascaro AL, Cesare P, Sacconi L, Grasselli G, Mandolesi G, MacO B, Knott GW, Huang L, De Paola V, Strata P, Pavone FS. 2013. In vivo single branch axotomy induces GAP-43-dependent sprouting and synaptic remodeling in cerebellar cortex. Proc Natl Acad Sci U S A 110:10824–10829. doi: 10.1073/pnas.1219256110

Antkowiak M, Torres-Mapa ML, Stevenson DJ, Dholakia K, Gunn-Moore FJ. 2013. Femtosecond optical transfection of individual mammalian cells. Nat Protoc 8:1216–1233. doi: 10.1038/nprot.2013.071

Barron HC, Vogels TP, Behrens TE, Ramaswami M. 2017. Inhibitory engrams in perception and memory. Proc Natl Acad Sci U S A. doi: 10.1073/pnas.1701812114

Bauer F, Michalowski A, Kiedrowski T, Nolte S. 2015. Heat accumulation in ultra-short pulsed scanning laser ablation of metals. Opt Express 23:1035. doi: 10.1364/oe.23.001035

Beiko D, Pierre SA, Leonard MP. 2011. Urethroscopic holmium:YAG laser epilation of urethral diverticular hair follicles following hypospadias repair. J Pediatr Urol 7:231–2. doi: 10.1016/j.jpurol.2010.09.018

Ben-David U, Benvenisty N. 2014. Chemical ablation of tumor-initiating human pluripotent stem cells. Nat Protoc 9:729–740. doi: 10.1038/nprot.2014.050

Blinder P, Friedman B, Shih AY, Tsai PS, Kleinfeld D, Stanley G, Lyden PD. 2012. The smallest stroke: occlusion of one penetrating vessel leads to infarction and a cognitive deficit. Nat Neurosci 16:55–63. doi: 10.1038/nn.3278

Bourgeois F, Ben-Yakar A. 2008. Femtosecond laser nanoaxotomy properties and their effect on axonal recovery in C. elegans: erratum. Opt Express 16:5963. doi: 10.1364/OE.16.005963

Buffelli M, Sacconi L, O’Connor RP, Pavone FS, Masi A, Jasaitis A. 2007. In vivo multiphoton nanosurgery on cortical neurons. J Biomed Opt 12:050502. doi: 10.1117/1.2798723

Cheng M, Qin X, Zhu F, Khalili K, Gordon J, Ohtake Y, Zhou Z, Peng X, Feng D, Yang X, Li S, Ju C, Zhang Y, Li M, Wang Hua, Zhao C, Gao B, He Y, Hayat U, Zhang L, Hu W, Wang Hong, Dai S, Kearns A, Xu M, Liu F, Bryda EC. 2016. Cre-inducible human CD59 mediates rapid cell ablation after intermedilysin administration. J Clin Invest 126:2321–2333. doi: 10.1172/jci84921

Chung SH, Mazur E. 2009. Femtosecond laser ablation of neurons in C. elegans for behavioral studies. Appl Phys A Mater Sci Process 96:335–341. doi: 10.1007/s00339-009-5201-7

Churgin MA, He L, Murray JI, Fang-Yen C. 2013. Efficient Single-Cell Transgene Induction in Caenorhabditis elegans Using a Pulsed Infrared Laser. Genes|Genomes|Genetics 3:1827–1832. doi: 10.1534/g3.113.007682

Cichon J, Gan W-B. 2015. Branch-specific dendritic Ca2+ spikes cause persistent synaptic plasticity. Nature 520:180–5. doi: 10.1038/nature14251

Damisah EC, Kwan AC, Grutzendler J, Hill RA, Chen F. 2017. Targeted two-photon chemical apoptotic ablation of defined cell types in vivo. Nat Commun 8:15837. doi: 10.1038/ncomms15837

Di Niso F, Gaudiuso C, Sibillano T, Mezzapesa FP, Ancona A, Lugarà PM. 2014. Role of heat accumulation on the incubation effect in multi-shot laser ablation of stainless steel at high repetition rates. Opt Express 22:12200. doi: 10.1364/OE.22.012200

Driscoll LN, Pettit NL, Minderer M, Chettih SN, Harvey CD. 2017. Dynamic Reorganization of Neuronal Activity Patterns in Parietal Cortex. Cell 170:986-999.e16. doi: 10.1016/j.cell.2017.07.021

Dunn FA. 2015a. Photoreceptor ablation initiates the immediate loss of glutamate receptors in postsynaptic bipolar cells in retina. J Neurosci 35:2423–2431. doi: 10.1523/JNEUROSCI.4284-14.2015

Dunn FA. 2015b. Photoreceptor Ablation Initiates the Immediate Loss of Glutamate Receptors in Postsynaptic Bipolar Cells in Retina 35:2423–2431. doi: 10.1523/JNEUROSCI.4284-14.2015

Eyo UB, Peng J, Swiatkowski P, Mukherjee A, Bispo A, Wu LJ. 2014. Neuronal hyperactivity recruits microglial processes via neuronal NMDA receptors and microglial P2Y12 receptors after status epilepticus. J Neurosci 34:10528–10540. doi: 10.1523/JNEUROSCI.0416-14.2014

Fino E, Yuste R. 2011. Dense inhibitory connectivity in neocortex. Neuron 69:1188–1203. doi: 10.1016/j.neuron.2011.02.025

Fouad AD, Teng S, Mark JR, Liu A, Alvarez-Illera P, Ji H, Du A, Bhirgoo PD, Cornblath E, Guan SA, Fang-Yen C. 2018. Distributed rhythm generators underlie Caenorhabditis elegans forward locomotion. Elife 7. doi: 10.7554/eLife.29913

Fu Y, Tucciarone JM, Espinosa JS, Sheng N, Darcy DP, Nicoll RA, Huang ZJ, Stryker MP. 2014. A cortical circuit for gain control by behavioral state. Cell 156:1139–1152. doi: 10.1016/j.cell.2014.01.050

Gabel C V., Antoine F, Chuang C-F, Samuel ADT, Chang C. 2008. Distinct cellular and molecular mechanisms mediate initial axon development and adult-stage axon regeneration in C. elegans. Development 135:1129–1136. doi: 10.1242/dev.030817

Gentet LJ, Kremer Y, Taniguchi H, Huang ZJ, Staiger JF, Petersen CC. 2012. Unique functional properties of somatostatin-expressing GABAergic neurons in mouse barrel cortex. Nat Neurosci 15:607–612. doi: 10.1038/nn.3051

Grégoire D, Kmita M. 2014. Genetic Cell Ablation. pp. 421–436. doi: 10.1007/978-1-60327-292-6_25

Guo SX, Bourgeois F, Chokshi T, Durr NJ, Hilliard MA, Chronis N, Ben-Yakar A. 2008. Femtosecond laser nanoaxotomy lab-on-a-chip for in vivo nerve regeneration studies. Nat Methods 5:531–533. doi: 10.1038/nmeth.1203

Hamad AH. 2016. Effects of Different Laser Pulse Regimes (Nanosecond, Picosecond and Femtosecond) on the Ablation of Materials for Production of Nanoparticles in Liquid SolutionHigh Energy and Short Pulse Lasers. InTech. p. 4203. doi: 10.5772/63892

Hayes JA, LaMar MD, Vann NC, Kottick A, Picardo MCD, Song H, Revill AL, Wang X, Akins VT, Del Negro CA, Funk GD. 2014. Laser ablation of Dbx1 neurons in the pre-Bötzinger complex stops inspiratory rhythm and impairs output in neonatal mice. Elife 3:1–25. doi: 10.7554/elife.03427

Hayes JA, Wang X, Del Negro CA. 2012. Cumulative lesioning of respiratory interneurons disrupts and precludes motor rhythms in vitro. Proc Natl Acad Sci 109:8286–8291. doi: 10.1073/pnas.1200912109

Hu W, Ferris SP, Tweten RK, Wu G, Radaeva S, Gao B, Bronson RT, Halperin JA, Qin X. 2008. Rapid conditional targeted ablation of cells expressing human CD59 in transgenic mice by intermedilysin. Nat Med 14:98–103. doi: 10.1038/nm1674

Ito M, Nagano N, Arai Y, Ogawa R, Kobayashi S, Motojima Y, Go H, Tamura M, Igarashi K, Dennery PA, Namba F. 2017. Genetic ablation of Bach1 gene enhances recovery from hyperoxic lung injury in newborn mice via transient upregulation of inflammatory genes. Pediatr Res 81:926–931. doi: 10.1038/pr.2017.17

Jasinski M. 2018. Numerical analysis of soft tissue damage process caused by laser action. AIP Conf Proc 1922. doi: 10.1063/1.5019063

Kobayashi J, Shidara H, Morisawa Y, Kawakami M, Tanahashi Y, Hotta K, Oka K. 2013. A method for selective ablation of neurons in C. elegans using the phototoxic fluorescent protein, KillerRed. Neurosci Lett 548:261–264. doi: 10.1016/j.neulet.2013.05.053

Krüger J, Kautek W. 2012. Ultrashort Pulse Laser Interaction with Dielectrics and Polymers. Springer, Berlin, Heidelberg. pp. 247–290. doi: 10.1007/b12683

Larkum M. E., Nevian T, Sandler M, Polsky A, Schiller J. 2009. Synaptic Integration in Tuft Dendrites of Layer 5 Pyramidal Neurons: A New Unifying Principle. Science (80-) 325:756–760. doi: 10.1126/science.1171958

Larkum Matthew E, Sandler M, Polsky A, Schiller J. 2009. Synaptic Integration in Tuft Dendrites of Layer 5 Pyramidal Neurons: A New Unifying Principle 663:756–761.

Latt SA, Stetten G, Juergens LA, Willard HF, Scher CD. 1975. Recent developments in the detection of deoxyribonucleic acid synthesis by 33258 Hoechst fluorescence. J Histochem Cytochem 23:493–505.

Li W, Ma L, Yang G, Gan WB. 2017. REM sleep selectively prunes and maintains new synapses in development and learning. Nat Neurosci 20:427–437. doi: 10.1038/nn.4479

Li X, Wu Z, Qin Z, He S, Wu W, Qu JY, Chen C, Sun Q, Luo Y, Lin Y. 2018. In vivo imaging-guided microsurgery based on femtosecond laser produced new fluorescent compounds in biological tissues. Biomed Opt Express 9:581. doi: 10.1364/boe.9.000581

Makhijani K, To TL, Ruiz-González R, Lafaye C, Royant A, Shu X. 2017. Precision Optogenetic Tool for Selective Single- and Multiple-Cell Ablation in a Live Animal Model System. Cell Chem Biol 24:110–119. doi: 10.1016/j.chembiol.2016.12.010

Mardinly AR, Oldenburg IA, Pégard NC, Sridharan S, Lyall EH, Chesnov K, Brohawn SG, Waller L, Adesnik H. 2018. Precise multimodal optical control of neural ensemble activity. Nat Neurosci 21:881–893. doi: 10.1038/s41593-018-0139-8

Morris RG, Amaral D, Bliss T, Keefe JO, Kropff E, Moss CF, Witter MP, Ulanovsky N, Nakamura K, Nishijo H, Eifuku S, Agnihotri NT, Streater S, Hawkins RD, Kandel ER, Mcnaughton BL, Poe GR, Collman F, Dombeck DA, Tank DW, Tang HM, Gohil BC, Botero JM, Kemere C, German PW, Frank LM, Verriotis MA, Jovalekic A, Fenton AA, Jeffery KJ, Anand RL, Anderson MI, Sirota A, Patel J, Cacucci F, Burgess N, Linder AN, Leutgeb JK, Leutgeb S, Tabuchi E, Matsumura N, Ono T, Shinder M, Derdikman D, Las L, Yovel Y, Ahrens M, Raz R, Kamenitz K, Pasmantirer B. 2013. Developmental Decline in Neuronal. Science (80-) 340:372–376.

Naylor RW, Chang HHG, Qubisi S, Davidson AJ. 2018. A novel mechanism of gland formation in zebrafish involving transdifferentiation of renal epithelial cells and live cell extrusion. Elife 7. doi: 10.7554/eLife.38911

Naylor RW, Dodd RC, Davidson AJ. 2016. Caudal migration and proliferation of renal progenitors regulates early nephron segment size in zebrafish. Sci Rep 6:1–14. doi: 10.1038/srep35647

Nimmerjahn A, Kirchhoff F, Kerr JND, Helmchen F. 2004. Sulforhodamine 101 as a specific marker of astroglia in the neocortex in vivo. Nat Methods 1:31–37. doi: 10.1038/nmeth706

Nishimura N, Schaffer CB, Friedman B, Lyden PD, Kleinfeld D. 2007. Penetrating arterioles are a bottleneck in the perfusion of neocortex. Proc Natl Acad Sci 104:365–370. doi: 10.1073/pnas.0609551104

Nishimura N, Schaffer CB, Friedman B, Tsai PS, Lyden PD, Kleinfeld D. 2006. Targeted insult to subsurface cortical blood vessels using ultrashort laser pulses: Three models of stroke. Nat Methods 3:99–108. doi: 10.1038/nmeth844

Orger MB, Kampff AR, Severi KE, Bollmann JH, Engert F. 2008. Control of visually guided behavior by distinct populations of spinal projection neurons. Nat Neurosci 11:327–333. doi: 10.1038/nn2048

Park J, Papoutsi A, Ash RT, Marin MA, Poirazi P, Smirnakis SM. 2019. Contribution of apical and basal dendrites to orientation encoding in mouse V1 L2/3 pyramidal neurons. Nat Commun 10. doi: 10.1038/s41467-019-13029-0

Pfeffer CK, Xue M, He M, Huang ZJ, Scanziani M. 2013. Inhibition of inhibition in visual cortex: The logic of connections between molecularly distinct interneurons. Nat Neurosci 16:1068–1076. doi: 10.1038/nn.3446

Pozner A, Xu B, Palumbos S, Gee JM, Tvrdik P, Capecchi MR. 2015. Intracellular calcium dynamics in cortical microglia responding to focal laser injury in the PC::G5-tdT reporter mouse. Front Mol Neurosci 8:1–10. doi: 10.3389/fnmol.2015.00012

Ralvenius WT, Haenraets K, Jegen M, Tudeau L, Haueter S, Conzelmann K-K, Johannssen H, Ghanem A, Hösli L, Foster E, Wildner H, Zeilhofer HU, Bösl M. 2015. Targeted Ablation, Silencing, and Activation Establish Glycinergic Dorsal Horn Neurons as Key Components of a Spinal Gate for Pain and Itch. Neuron 85:1289–1304. doi: 10.1016/j.neuron.2015.02.028

Rompolas P, Greco V, Saotome I, Zito G, Haberman AM, Deschene ER, Gonzalez DG. 2012. Live imaging of stem cell and progeny behaviour in physiological hair-follicle regeneration. Nature 487:496–499. doi: 10.1038/nature11218

Saito M, Iwawaki T, Taya C, Yonekawa H, Noda M, Inui Y, Mekada E, Kimata Y, Tsuru A, Kohno K. 2001. Diphtheria toxin receptor-mediated conditional and targeted cell ablation in transgenic mice. Nat Biotechnol 19:746–750. doi: 10.1038/90795

Sandler M, Shulman Y, Schiller J. 2016. A Novel Form of Local Plasticity in Tuft Dendrites of Neocortical Somatosensory Layer 5 Pyramidal Neurons. Neuron 90:1028–1042. doi: 10.1016/j.neuron.2016.04.032

Strickland D, Mourou G. 1985. Compression of amplified chirped optical pulses. Opt Commun 55:447–449. doi: 10.1016/0030-4018(85)90151-8

Tertilt C, Heinen TJAJ, Wunderlich FT, Waisman A, Heppner FL, Jung S, Buch T, Kremer M. 2005. A Cre-inducible diphtheria toxin receptor mediates cell lineage ablation after toxin administration. Nat Methods 2:419–426. doi: 10.1038/nmeth762

Tsai PS, Blinder P, Migliori BJ, Neev J, Jin Y, Squier JA, Kleinfeld D. 2009. Plasma-mediated ablation: an optical tool for submicrometer surgery on neuronal and vascular systems. Curr Opin Biotechnol 20:90–99. doi: 10.1016/j.copbio.2009.02.003

Tsai PS, Friedman B, Ifarraguerri AI, Thompson BD, Lev-Ram V, Schaffer CB, Xiong Q, Tsien RY, Squier JA, Kleinfeld D. 2003. All-optical histology using ultrashort laser pulses. Neuron 39:27–41. doi: 10.1016/S0896-6273(03)00370-2

Urban-Ciecko J, Barth AL. 2016. Somatostatin-expressing neurons in cortical networks. Nat Rev Neurosci 17:401–409. doi: 10.1038/nrn.2016.53

Vogel A, Noack J, Hüttman G, Paltauf G. 2005. Mechanisms of femtosecond laser nanosurgery of cells and tissues. Appl Phys B Lasers Opt 81:1015–1047. doi: 10.1007/s00340-005-2036-6

Wang X, Hayes JA, Picardo MCD, Del Negro CA. 2013. Automated cell-specific laser detection and ablation of neural circuits in neonatal brain tissue. J Physiol 591:2393–2401. doi: 10.1113/jphysiol.2012.247338

Watanabe W, Arakawa N, Matsunaga S, Higashi T, Fukui K. 2004. Femtosecond laser disruption of subcellular organelles in a living cell 12:113–115.

Wilde GJC, Sundström LE, Iannotti F. 1994. Propidium iodide in vivo: an early marker of neuronal damage in rat hippocampus. Neurosci Lett 180:223–226. doi: 10.1016/0304-3940(94)90525-8

Wu Z, Ghosh-Roy A, Yanik MF, Zhang JZ, Jin Y, Chisholm AD. 2007. Caenorhabditis elegans neuronal regeneration is influenced by life stage, ephrin signaling, and synaptic branching. Proc Natl Acad Sci 104:15132–15137. doi: 10.1073/PNAS.0707001104

Xu H, Jeong HY, Tremblay R, Rudy B. 2013. Neocortical Somatostatin-Expressing GABAergic Interneurons Disinhibit the Thalamorecipient Layer 4. Neuron 77:155–167. doi: 10.1016/j.neuron.2012.11.004

Yanik MF, Cinar H, Cinar HN, Chisholm AD, Jin Y, Ben-Yakar A. 2004. Functional regeneration after laser axotomy. Nature 432:822. doi: 10.1038/432822a

